# Parvalbumin protein controls inhibitory tone in the spinal cord

**DOI:** 10.1101/2022.09.15.508019

**Authors:** Haoyi Qiu, Lois Miraucourt, Hugues Petitjean, Albena Davidova, Philipa Levesque-Damphousse, Jennifer L. Estall, Reza Sharif-Naeini

## Abstract

The nervous system processes sensory information by relying on the precise coordination of neuronal networks and their specific synaptic firing patterns. In the spinal cord, disturbances to the firing pattern of the tonic firing parvalbumin (PV)-expressing inhibitory interneuron (PV neurons) disrupt the ability of the dorsal horn to integrate touch information and may result in pathological phenotypes. The parvalbumin protein (PVp) is a calcium (Ca^2+^)-binding protein that buffers the accumulation of Ca^2+^ following a train of action potential to allow for tonic firing. Here, we find that peripheral nerve injury causes a decrease in PVp expression in PV neurons and makes them transition from tonic to adaptive firing. We also show that reducing the expression of PVp causes otherwise healthy adult mice to develop mechanical allodynia and causes their PV neurons to lose their high frequency firing pattern. We show that this frequency adaptation is mediated by activation of SK channels on PV neurons. Further, we show their tonic firing can be partially restored after nerve injury by selectively inhibiting the SK2 channels of PV neurons. We also reveal that a decrease in the transcriptional coactivator, PGC-1α, causes decrease PVp expression and the development of mechanical allodynia. By preventing the decrease in PVp expression before nerve injury, we were able to protect mice from developing mechanical allodynia. Our results indicate an essential role for PVp-mediated calcium buffering in PV neuron firing activity and the development of mechanical allodynia after nerve injury.

## Introduction

The intricate function of the nervous system relies on the coordination of neuronal networks that work together to process information, interpret sensations, and generate behavior. Each neuron encodes signals using specific firing frequencies that are finely tuned to orchestrate complex behaviors within the system, akin to a symphony where the proper function and timing of every instrument is critical to the ensemble. The basis for these precise firing rates relies on individual ionic conductances and their regulatory mechanisms.

In many areas of the central nervous system (CNS), one essential component of neuronal circuits is the tonic firing inhibitory interneuron (Bartos, Vida & Jonas, 2007; Rudy et al., 2011; Hughes & Todd, 2020). Whether they are in the hippocampus, the striatum, or the dorsal horn of the spinal cord, these tonic firing inhibitory neurons control the output of the circuits in which they are embedded (Amilhon et al., 2015; Fuchs et al., 2007; Espinoza et al., 2018; Orduz et al., 2013; Petitjean et al., 2015; Boyle et al., 2019). In the dorsal horn of the spinal cord, these inhibitory neurons control the outflow of nociceptive (painful) information from the spinal cord to the brain (Hughes et al., 2012; Foster et al., 2015; Petitjean et al., 2015; Boyle et al., 2019; Gradwell et al., 2022). Indeed, in the dorsal horn, these tonic firing inhibitory neurons function as gate keepers of touch-evoked pain sensation by preventing peripheral touch inputs from activating nociceptive circuits (Petitjean et al., 2015). A common feature of these tonic firing inhibitory neurons across all regions of the central nervous system is that they can relay critical information through the precise pattern of their actional potentials (Hughes et al., 2012; Amilhon et al., 2015; Espinoza et al., 2018; Boyle et al., 2019). Therefore, even the slightest disturbance to their firing pattern may impair circuit function and be associated with pathological phenotypes.

The current understanding of the mechanisms responsible for this tonic firing pattern is a dependence on the tight regulation of intracellular calcium levels (Zündorf & Reiser, 2011; Albéri et al., 2013; Orduz et al., 2013). Indeed, it has been reported that artificially manipulating intracellular calcium concentration can change the tonic firing pattern of these neurons, presumably through the recruitment of Ca^2+^-activated potassium channels of the small conductance (SK) family (Bischop et al., 2012; Orduz et al., 2013). These observations indicate that tonic firing neurons likely possess endogenous mechanisms that buffer cytoplasmic calcium ions and enable high-frequency firing. Identifying these calcium buffering processes is critical to understanding the firing pattern of inhibitory neurons and may shed light on pathologies associated with defective inhibitory tone.

One endogenous calcium buffer that is common to fast spiking neurons across the CNS is the parvalbumin protein (PVp), a calcium buffer containing two functional EF-hand type calcium binding sites (Kretsinger & Nockolds, 1973; Kawaguchi et al., 1987; Lewit-Bentley & Rety, 2000). Evidence for the correlation between PVp and PV neuron excitability is demonstrated in hippocampal neurons, where the activation of PV neurons induced high PVp expression while their inhibition induced a low PVp expression (Donato, Rompani & Caroni, 2013). In the striatum, perforated patch recordings demonstrated that knocking out PVp impacted the rhythmicity of PV neuron spiking (Orduz et al., 2013). Whether a similar link between PVp expression and PV neuron activity is observed in the spinal cord is unknown. This link is especially relevant because recent reports indicate that following nerve injury and the development of chronic pain, the excitability of PV neurons is decreased (Boyle et al., 2019). However, whether this change in firing pattern is caused by alterations to PVp remains to be determined.

In this study, we investigated the role of PVp in enabling the tonic firing of PV neurons, and how changes in the expression of this chelator affect somatosensory processing. We show that after nerve injury, spinal cord PV neurons change their firing pattern and start displaying frequency adaptation. This is correlated temporally with a decrease in PVp expression and the presence of mechanical allodynia, a painful response to an innocuous touch-like stimulus. To examine the causal relation between these events, we reduced PVp expression in healthy mice and observed a change in firing pattern as well as the development of mechanical allodynia. Finally, we demonstrate that overexpressing PVp in PV neurons prior to nerve injury attenuates the development of mechanical allodynia. These findings demonstrate the critical role of the parvalbumin protein in controlling the electrical activity of these inhibitory interneurons and in the preventing touch inputs from activating central pain circuits.

## Results

### The majority of dorsal horn PV neurons are inhibitory

Unlike in the cortex, where PV neurons are inhibitory, spinal PV neurons have been shown to be either excitatory or inhibitory, with the ratios ranging from 50 to 95% inhibitory (Hughes et al., 2012; Petitjean et al., 2015; Abraira et al., 2017; Gradwell et al., 2022). Using a multi-labelling fluorescence *in situ* hybridization approach on C57BL/6 male mice, we examined the neurochemical nature of the lamina IIi-III dorsal horn PV neurons. Our experiments revealed that 65.2 ± 3.9% (mean ± standard error of the mean, s.e.m.) of neurons that express the Pvalb mRNA also express the Slc32a1 mRNA which encodes the vesicular GABA transporter (VGAT), a specific marker of inhibitory interneurons (Sagné et al. 1997; Bedet et al. 2000) **(Figure 1A-C)**. This expression was at a higher intensity than Slc17a6 mRNA which encodes the vesicular glutamatergic transporter type 2 (VGLUT2), a specific marker of excitatory interneurons in the dorsal horn (Todd et al. 2003; Levine et al. 2014) **(Figure S1A-C)**. Furthermore, 34.9 ± 3.9% (mean ± s.e.m.) of the Pvalb mRNA expressing cells expressed the converse, higher intensity of VGLUT2 than VGAT indicating these neurons were excitatory.

**Figure 1.**
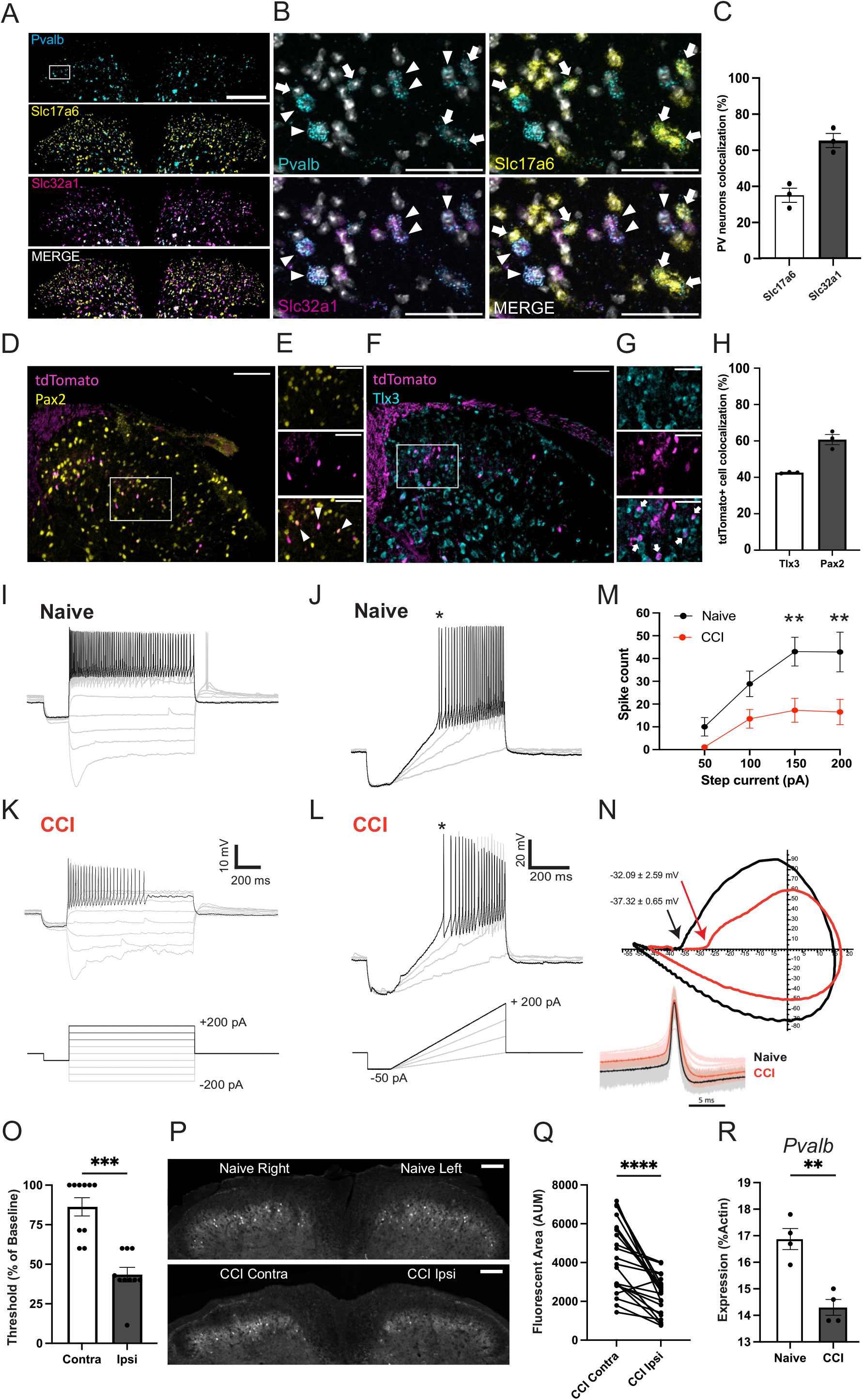
Dorsal horn PV neurons are primarily inhibitory interneurons and decrease their parvalbumin expression and firing frequency after nerve injury. **A.** Fluorescent *in situ* hybridization of mouse spinal cord dorsal horn reveals that neurons containing parvalbumin mRNA (cyan) expresses both VGAT mRNA (Slc32a1; inhibitory marker, magenta) and VGLUT2 mRNA (Slc17a6; excitatory marker, yellow). Scale bar, 200μm. **B.** Zoomed in image of A white box. White arrows show colocalization of Pvalb and Slc17a6 indicating excitatory PV interneurons. White arrowheads show colocalization of Pvalb and Slc32a1 indicating inhibitory PV interneurons. NucBlue is shown in grey. Scale bar, 50 μm. **C.** Quantification of the mean percentage of colocalization (±SEM) between PV mRNA and inhibitory VGAT mRNA (black, 902/1390 PV+ cells, n=33 sections from 3 mice) and between PV mRNA and excitatory VGLUT2 mRNA (white, 488/1390 PV+ cells, n=33 sections from 3 mice). **D.** Transverse sections of the dorsal horn neurons stained with Pax2 antibodies (yellow). tdTomato-positive neurons are shown in magenta. Scale bar, 100μm. **E.** Zoomed in image of D white box. Top panel shows Pax2 staining, middle panel shows tdTom-positive PV neurons. Bottom panel shows merged image. White arrowheads show colocalization of Pax2 and tdTomato. Scale bar, 50μm. **F.** Transverse sections of the DH neurons stained with Tlx3 antibodies (cyan) tdTomato-positive PV neurons in magenta. Scale bar, 100μm. **G.** Zoomed in image of F white box. Top panel shows Tlx3 staining, middle panel shows tdTom-positive PV neurons. Bottom panel shows merged image. White arrowheads show colocalization of Tlx3 and tdTomato. Scale bar, 50μm. **H.** Most tdTom-positive PV neurons colocalize with Pax2 (black, 60.76 ± 2.8% counted 319 cells from 3 mice), while less tdTom-positive PV neurons colocalize with Tlx3 (white, 42.61 ± 2.4% counted from 227 cells from 3 mice). **I-L** Whole cell current clamp raw traces illustrating the response to step or ramp current injections of PV neurons recorded in lumbar dorsal horn spinal cord of naïve and nerved injured (CCI) mice. **M.** Spike count analysis (n=5 cells from 3 naïve mice, n=8 cells from 4 CCI mice), mean ± S.E.M, **p<0.01 Two-way ANOVA, Šídák's multiple comparisons test. **N.** The spike threshold for each PV neuron was calculated based on the first spikes (*) evoked on depolarizing ramp, which were then averaged for all the “Naïve” (black) or “CCI” (red) recordings and plotted using the first derivative of the voltage (dV/dt) against the voltage (V). **O.** Mechanical threshold (percent of baseline) of nerve injured animals decreases in the ipsilateral hind paw compared to the contralateral paw (n=10 mice) ***p-value=0.0008, paired t-test. **P.** Transverse lumbar sections of nerve injured (CCI) Pvalb-tdTomato mice showing a decrease in Pvalb promoter activity (decrease in tdTom+ cells) on the ipsilateral side only. Scale bar, 100μm. **Q.** There is a decrease in the area of tdTom+ cells per spinal cord section (n=22 sections in 2 CCI mice) ****p-value <0.0001, paired t-test. No changes were observed in the area of tdTom+ cells of naïve mice (n=20 sections in 2 mice; data not shown). **R.** Mean (±SEM) of Pvalb mRNA normalized to actin in the dorsal ipsilateral quadrant of naïve and CCI mice (n=4 mice per group). There is also an upregulation of PKCγ, AIF1, and GFAP in the CCI mice used as positive controls (data not shown). **p value=0.0021, unpaired t-test.

To determine whether these distributions of inhibitory and excitatory PV neurons were maintained at the protein level, we performed immunohistochemistry experiments with antibodies against Pax2 and Tlx3, markers of excitatory and inhibitory neurons, respectively (Gross et al. 2002; Cheng et al. 2004). These experiments were performed in mice generated by crossing PV^cre^ mice (JAX, stock #017320) to the Ai14 tdTomato reporter (JAX, stock#007914), referred to as PV^cre^;tdTom mice. Our results indicate that these heterozygous PV^cre^;tdTom mice were only able to capture 33.8 ± 2.2% of PV-immunoreactive (IR) neurons in the lumbar dorsal horn, yet they were highly specific (81.5 ± 2.3%) (**Figure S1D-E**). These proportions are similar to those reported in a recent study (Gradwell et al., 2022). Furthermore, our results indicate that in lamina IIi-III of the dorsal horn, 60.8 ± 2.8% of tdTom-positive neurons colocalized with PAX2 whereas only 42.61 ± 2.4% colocalized with Tlx3 (**Figure 1D-H**). Taken together, this data shows that the majority of lamina IIi-III dorsal horn PV interneurons are inhibitory interneurons.

### PV neurons lose their ability to tonically fire after nerve injury

We previously showed that inhibitory PV neurons function as modality-specific filters of sensory inputs in the dorsal horn. Silencing their activity in naïve mice induces mechanical allodynia. On the other hand, selectively activating them in a mouse model of neuropathic pain alleviates mechanical allodynia (Petitjean et al., 2015). After nerve injury, there is a loss of dorsal horn inhibitory tone driven partly by a decrease in the function, but not in the number, of inhibitory PV neurons (Petitjean et al., 2015; Boyle et al., 2019). We hypothesized that this decreased function can be the result of decreased appositions onto their targets, reduced excitatory drive from Aβ fibers, or altered electrical activity of PV neurons. We therefore examined the electrical properties of PV neurons after nerve injury induced by chronic constriction to the sciatic nerve (CCI). When examined at 4 weeks post-CCI, a time point when mice display significant mechanical allodynia (**Fig 1O**), PV neurons lose their ability to fire tonically. Indeed, we demonstrate that PV neurons display frequency adaptation and can no longer sustain high frequency firing patterns in response to current step injections **(Figure 1I, K, L)** and to depolarizing current ramp **(Figure 1J, M**). However, we failed to observe any change in their resting membrane potential after nerve injury (−65.26 ± 0.70 mV in naïve versus −60.73 ± 1.96 mV in CCI, p=0.0597 unpaired t-test). We also found no statistically significant differences in the action potential threshold (−37.32 ± 0.65 mV naïve versus −32.09 ± 2.59 mV CCI, p=0.0892 unpaired t-test) (**Figure 1N**). Our data indicate that after nerve injury, PV neurons change their action potential firing pattern from tonic to adaptive firing, which likely causes a decrease in their inhibitory output.

### PV expression decreases after nerve injury

It has been shown that striatal PV neuron excitability is linked to the calcium buffering capacity of the cell (Orduz et al., 2013). One of the most highly expressed calcium buffers in PV neurons is the PV protein itself. To examine whether nerve injury leads to a change in PVp expression, we used a BAC transgenic mouse in which the PV promoter drives the expression of tdTomato (Pvalb-tdTomato mice) (Meyer et al. 2002). To focus our analysis on the PV neurons located in the spinal segment affected by the nerve injury, we narrowed our quantification to sections displaying increased microglia staining on the ipsilateral side compared to the contralateral side (**Figure S1F-G**). This approach is based on the observation that nerve injury causes an increase in the proliferation of microglia in the dorsal horn segment receiving inputs from the injured area (Piao et al. 2006). The expression of tdTomato is distributed throughout the soma and processes of these PV neurons. Therefore, in a region of interest encompassing lamina IIi and III, we quantified the area of tdTomato fluorescence. Our results show that in this region, 4 weeks after CCI, the dorsal horn displays a significantly lower area of tdTomato fluorescence ipsilateral to the injury compared to the contralateral side (**Figure 1P-Q**) or to naïve mice (data not shown). Furthermore, we observed a decrease in the tdTomato intensity of individual cells in nerve injured animals compared to naïve, and in the ipsilateral compared to the contralateral side, indicating a decrease in PV promoter activity after nerve injury (**Figure S1H**). These findings were supported by real time (RT)-qPCR experiments performed on ipsilateral and contralateral dorsal horn quadrants of CCI and naïve mice, indicating a decrease in *Pvalb* mRNA **(Figure 1R**). To ensure the selected quadrants represented segments innervated by the sciatic nerve, we confirmed the increase of other mRNAs known to be upregulated after nerve injury, such as AIF1, GFAP and PKCγ (Mao, et al., 1995; De Luca et al., 2022). Our results show that 4 weeks after peripheral nerve injury, PV neurons have significantly less PV expression, which may impact the cell’s calcium handling capacity and calcium-dependent conductances, eventually affecting the neurons’ firing properties and their inhibitory output.

### Decreasing PV protein leads to the development of mechanical allodynia

Although mechanical allodynia and the decrease in PVp are both observed after nerve injury, the presence of a causal relation between the two remained unknown. Indeed, nerve injury recruits several inflammatory processes in the dorsal horn, making it difficult to ascribe a role to the decrease in PV protein expression to the development of mechanical allodynia. Therefore, we examined whether deleting PV protein would be enough to elicit the development of mechanical allodynia. We assessed the sensitivity to thermal and mechanical stimuli in PV knockout mice (PV KO mice, JAX stock #010777). Compared to wild type mice, PV KO mice displayed similar heat sensitivity (**Figure 2A**) but were hypersensitive to static mechanical stimuli akin to what is observed in mice following nerve injury (**Figure 2A, B**). These findings confirmed that PV neurons are modality specific filters of mechanical inputs (Petitjean et al., 2015). However, results obtained from global KO mice may be confounded by the possible developmental compensation and the impact of the gene deletion in other CNS areas implicated in pain transmission or interpretation. Therefore, we sought to reduce PV protein expression in otherwise healthy adult wild type mice and examine their sensitivity to mechanical stimuli. Wild type Pvalb-tdTomato mice were injected with lentiviruses containing shRNA molecules against PV (shPV) or control non-targeting (NT) sequences. We confirmed that PVp was decreased in shPV-injected mice by quantifying the tdTomato fluorescence intensity around the injection site (**Figure S2A, S2B**). Our results indicated that mice injected with the shPV virus developed mechanical allodynia, but not those injected with the NT virus (**Figure 2C, S2D**). This increased sensitivity was modality-specific because thermal sensitivity was unaffected (**Figure S2C**). These results indicate that a reduction in PV protein expression alone is enough to induce mechanical allodynia similar to that seen after nerve injury and after silencing PV neurons (Petitjean et al., 2015).

**Figure 2.**
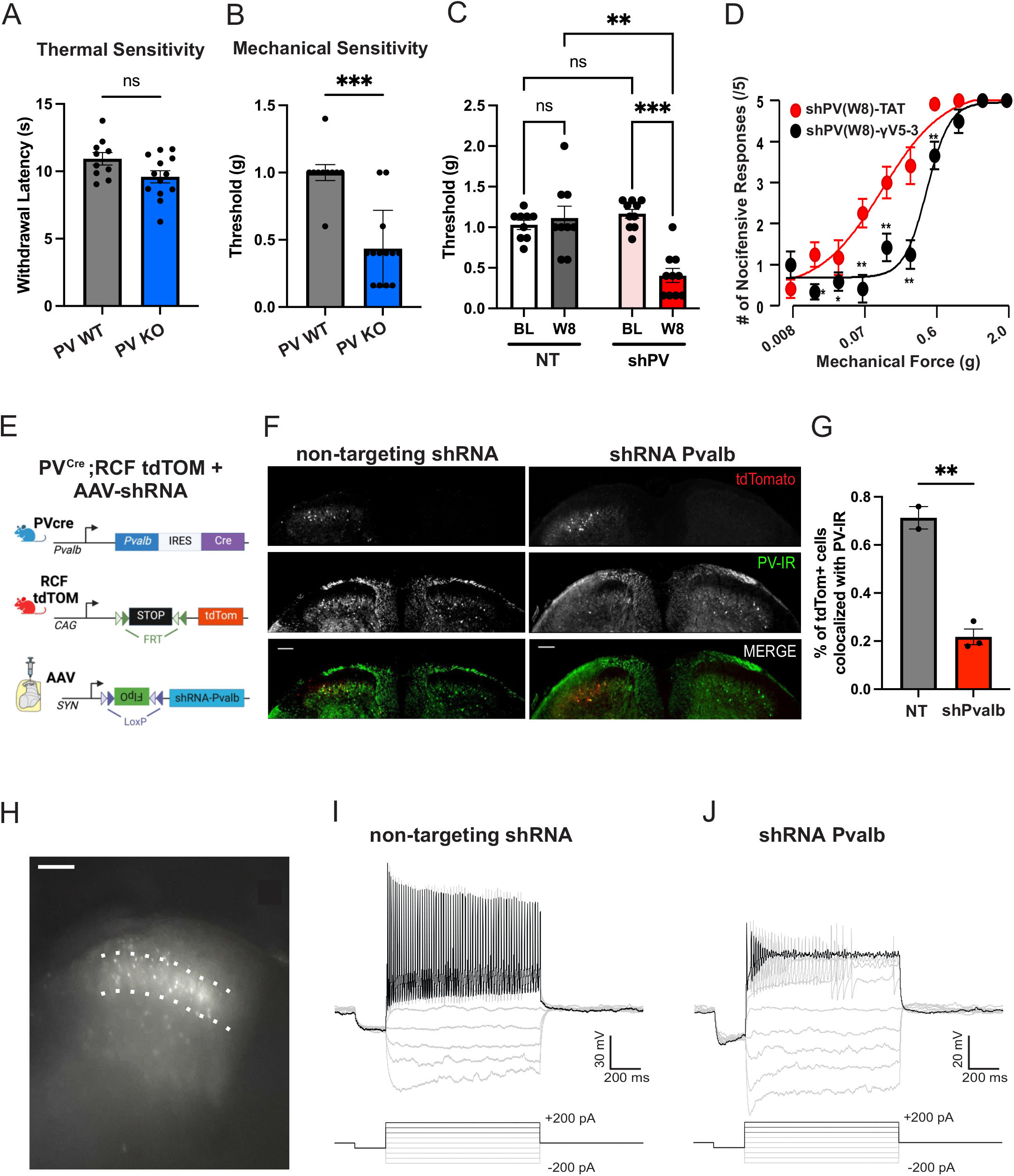
Decreasing PV protein in naïve mice elicits mechanical hypersensitivity and decreases firing frequency of PV neurons. **A.** Compared to PV wild type (PV WT) mice (grey bar), PV knockout (PV KO) mice (blue bar) show similar paw withdrawal latencies to heat (n=13 KO mice, n=11 WT mice, ns. not statistically significant, unpaired t-test). **B.** PV KO mice (blue bar) show decreased pain thresholds to mechanical stimuli compared to PV WT mice (grey bar) (***p-value<0.001, unpaired t-test). **C.** Intraspinal delivery of a lentivirus carrying shRNA against PV (n =10 mice, pink and red bars) produces mechanical allodynia after 8 weeks (W8) compared to baseline (BL). Mice receiving scrambled shRNA (n =9 mice, white and black bars) are unaffected. Two-way ANOVA, **p-value=0.0088, ***p-value<0.001, Šídák's multiple comparisons test. **D.** Mean (±SEM) mechanically evoked (von Frey) nocifensive responses in shRNA mice injected against PV after 8 weeks post injection (shPV-W8) 20 minutes after receiving an intrathecal injection of the control TAT peptide (red dots and curve, control) state the presence of a mechanical allodynia. Following the injection of the inhibitor of the PKCγ enzyme, γV5-3, there is a significant shift in the mechanical sensitivity (black dots and curve) (n=6 mice shPV-W8-TAT, n=6 mice shPV-W8-γV5-3). Two-way ANOVA, Tukey’s multiple comparisons test. **E.** Schematic diagram of experimental design to visualize shRNA-Pvalb expressing PV neurons for spinal cord slice electrophysiology. A PV^cre^ mouse expressing Cre recombinase in Pvalb-expressing neurons (top row) is crossed with an RCF-tdTomato mouse expressing *frt-*flanked STOP cassette preventing the transcription of tdTomato fluorescent protein (middle row). The offspring is intraspinally injected with an AAV containing a cre-dependent FlpO shRNA-Pvalb. Thus, only the PV neurons expressing the AAV will be tdTomato+ and recorded from. **F-G.** Transverse dorsal horn sections of PV^cre^;RCF tdTOM mice injected with either non-targeting/scramble shRNA (left) or shRNA-Pvalb (right). Anti-PV staining (green) showed that 21.79 ± 3.3% of tdTomato+ cells were PV-IR+ in the shRNA-Pvalb injected animals (n=21 sections from 3 mice, red bar), whereas 71.33 ± 4.7% of tdTomato+ cells were PV-IR+ in the non-targeting control (n=13 sections from 2 mice, grey bar). **p-value = 0.0028, unpaired t-test. Scale bar, 100μm. **H.** Microphotography of a lumbar spinal cord slice obtained from a shRNA-Pvalb injected PV^cre^;RCF tdTom mouse showing tdTomato+ under an epifluorescence microscope used for targeted electrophysiology recordings. Scale bar, 100μm. **I, J.** Whole cell current clamp raw traces illustrating the response to incremental current step injections of a tdTomato+ non-targeting shRNA and shRNA-Pvalb expressing PV neuron recorded in lumbar dorsal horn spinal cord.

### The allodynic effect of decreasing PVp expression is alleviated by PKCγ inhibition

We postulate that the mechanical allodynia induced by decreasing PV protein is through the disinhibition of the postsynaptic targets of PV neurons. Blocking the activity of these postsynaptic targets should therefore alleviate the allodynia induced by the decrease in PVp expression. One of the targets of PV neurons are PKCγ-expressing excitatory neurons (PKCγ neurons) (Petitjean et al. 2015, Boyle et al., 2019). To examine the proportion of PKCγ neurons contacted by PV neurons, we used a cre-dependent anterograde transsynaptic tracer - wheat germ agglutin (WGA) – packaged in an AAV vector (Peirs et al. 2015; François et al. 2017) and injected in PV^cre^;tdTom mice (Figure S2E). Under these conditions, PV neurons were the source of WGA production and transferred it to their post-synaptic targets. When examined two weeks after virus injection, immunohistochemistry experiments revealed the presence of WGA in several non-PV neurons, 17.6% of which were PKCγ neurons (**Figure S2E-S2F**). Importantly, nearly 80% of WGA-IR neurons were not labeled by PKCγ, indicating that PV neurons may have several other targets in the dorsal horn, including the recently reported vertical cells (Boyle et al., 2019). It was recently reported that the inhibitory output of PV neurons onto PKCγ neurons was significantly lower in nerve injured mice (Chen, Scherrer & Cao, 2022), further supporting the notion of a disinhibition of postsynaptic targets. In these excitatory neurons, activity of the PKCγ enzyme was shown to be central to their role in mechanical allodynia (Malmberg et al., 1989; Miraucourt, Dallel & Voisin 2007; Petitjean et al., 2015). To examine whether a disinhibition of PKCγ neurons was driving the mechanical allodynia elicited by the downregulation of PVp, we administered an intrathecal injection of the PKCγ blocker γV5-3 (or the control TAT peptide) 8 weeks after shPV injection. Our results revealed that inhibiting PKCγ in these mice significantly alleviated their mechanical allodynia (**Figure 2D**). Taken together, our results showed that reducing PVp expression in an otherwise healthy mouse precipitates a mechanical allodynia akin to nerve injury, which is mediated in part by the disinhibition of PKCγ neurons.

### Decreasing PV protein impairs the ability of PV neurons to fire tonically

We anticipated that if decreasing the expression of a calcium chelator (PVp) causes mechanical allodynia, it would be through the impairment of the electrical activity of these neurons. Thus, we examined if decreasing PVp expression in otherwise healthy animals perturbs the PV neuron firing pattern, similar to what is observed after nerve injury. To address this question, we developed a new mouse strain and used an AAV vector (**Figure 2E**) to visually identify PV neurons with downregulated PVp expression for spinal cord slice electrophysiology (**Figure 2H**). We verified the decrease in PV-immunoreactivity of the tdTomato+ cells and the development of mechanical allodynia in mice that received shPvalb compared to non-targeting shRNA 10 days after injection (**Figure 2F-G, S2G**). Furthermore, the tdTomato+ cells of shRNA-Pvalb injected mice showed decreased firing activity compared to the non-targeting control (**Figure 2I-J**). We should note that injection of the non-targeting shRNA produced a mild allodynia which may be attributed to the intraspinal injection. Taken together, our results indicate that a decrease in PVp alone is enough to trigger the development of mechanical allodynia and change the firing activity of PV neurons.

### Active inhibition of SK channels enables the tonic firing ability of PV neurons

We showed that the firing pattern of PV neurons changes after nerve injury. The disruption in the tonic firing could be the result of a decrease in conductances that enable tonic firing, such as HCN channels (Roth & Hu, 2020), or an increase in conductances that oppose tonic firing, such as SK channels (Orduz et al., 2013). Previous studies showed inhibitory PV neuron of dorsal horn lamina IIi-III highly express HCN4 channels responsible for establishing their hyperpolarization-activated (Ih) current (Hughes et al., 2012, Hughes et al., 2013). HCN channels are known to enhance action potential initiation during tonic firing and facilitate the propagation of action potentials in PV expressing basket cells of the hippocampus (Roth & Hu, 2020). Furthermore, blocking the Ih current with the pan-selective antagonist ZD7288 in neuropathic pain models provided significant analgesic effects (Chaplan et al., 2003; Tu et al, 2006). Thus, we examined whether their inhibition in naïve mice could impair tonic firing activity of PV neurons. Our results show that bath application of ZD7288 does not significantly reduce the firing frequency of dorsal horn PV neurons (**Figure S3A-C**) or impact the resting membrane potential (**Figure S3D**). This inability of HCN blockade to impact tonic firing has also been observed in the hippocampus (Aponte et al., 2006; Roth & Hu, 2020). These observations indicate that blocking HCN channels does not cause a change in the firing properties of PV neurons.

Another important contributor to the firing pattern of PV neurons are the calcium-activated SK channels (Orduz et al., 2013). We hypothesized that the nerve injury induced decrease in PVp expression may reduce the ability of the PV neuron to chelate action potential-driven calcium influx, thus enabling the activation of SK channels and the shift from tonic to adaptive firing patterns. To determine whether PV neurons express SK channels, we examined the expression of the SK channel isoforms (Kcnn1-4) using RT-qPCR on spinal dorsal quadrants and found that all four isoforms were present (**Figure 3A**). To determine whether these channels play a functional role, we examined the effect of their activation on the tonic firing activity of PV neurons. Our results demonstrated that the application of the SK channel agonist, 1-EBIO, can reversibly impose frequency adaptation in tonic firing PV neurons **(Figure 3B-D)**. Because SK2 (Kcnn2) and SK3 (Kcnn3) channels were more highly expressed in our RT-qPCR results of spinal dorsal quadrants, we focused our efforts on these channels. This observation is supported by a recently published dorsal horn single cell RNA-sequencing database (Häring et al., 2018), reporting high expression of Kcnn2 and low expression of Kcnn3 in the same inhibitory neuronal cluster as Pvalb. We further confirmed the expression of these channel isoforms by *in situ* hybridization and demonstrated that Kcnn2 is the predominant isoform of SK channels in dorsal horn inhibitory PV neurons (**Figure 3E, H**).

**Figure 3.**
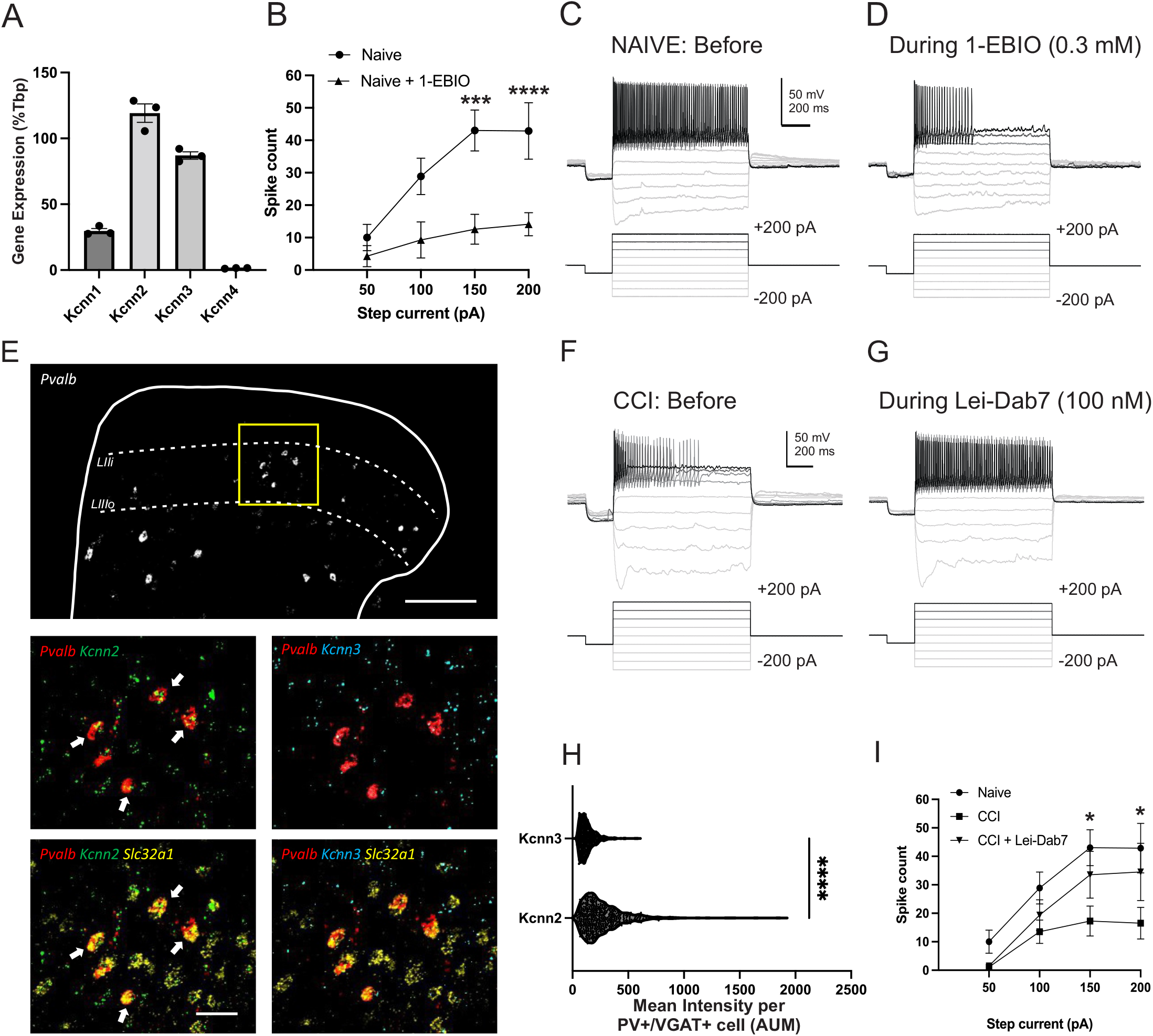
Inhibitory PV neurons express functional SK2 channels responsible for frequency adaptation after nerve injury. **A.** qPCR of lumbar spinal dorsal quadrants of Kcnn1 (29.67 ± 1.9%), Kcnn2 (119.2 ± 7.0%), Kcnn3 (87.0 ± 2.8%), Kcnn4 (1.33 ± 0.16%) normalized to the control gene, TATA box binding protein (Tbp) (n=3 mice). **B-D.** Whole cell current clamp raw traces illustrating the response to current step injections of a PV neuron recorded in lumbar spinal cord slices of a naïve mouse, before and after bath application of the SK channel activator 1-EBIO (0.3 mM) and the spike count analysis (mean ± S.E.M, n=7 cells from 3 mice). * p<0.05, **p<0.01 Two-way ANOVA, Bonferroni's multiple comparisons test. **E**. Fluorescence *in situ* hybridization of transverse lumbar spinal dorsal horn sections (white outline) showing PV mRNA (red), VGAT mRNA (slc32a1, yellow), SK2 mRNA (kcnn2, green) and SK3 mRNA (kcnn3, cyan) colocalization. Zoom in sections (yellow box) shows white arrows indicating inhibitory PV neurons colocalizing with kcnn2. Scale bar, 100μm, 50μm. **F-G.** Whole cell current clamp raw traces illustrating the response to incremental current step injections of a PV neuron recorded in lumbar spinal cord slices of a CCI mouse, before and after bath application of the SK2 channel specific blocker Lei Dab-7 (0.1 nM). **H.** Quantification of E indicating inhibitory PV neurons have a significantly higher expression of Kcnn2 [273.5 ± 7.745 Intensity arbitrary units of measurements (AUM)] than Kcnn3 (130.9±2.722 Intensity AUM). Total of 684 VGAT+/PV+ cells counted in 3 mice, ****p-value <0.0001 paired t-test. **I.** The spike count analysis of PVINs from naïve, CCI and CCI with bath application of Lei-Dab7. (mean ± S.E.M, n=8 cells from 4 mice CCI, n=7 cells from 3 mice Naive). Two-way ANOVA, Šídák's multiple comparisons test. *p<0.01 compares Naïve and CCI. No statistically significant difference between naïve and CCI+Lei-Dab7.

We hypothesize that after nerve injury, the decrease in PVp expression enables the accumulation of intracellular calcium, which activates SK2 channels to produce frequency adaptation. Consequently, inhibiting SK2 channels in PV neurons of nerve-injured mice should reverse their adaptive behavior and restore tonic firing. Indeed, our results demonstrated that bath application of Lei-Dab7, a selective blocker of SK2 channels (Shakkottai et al., 2001), partially restored the tonic firing of CCI PV neurons **(Figure 3F-G, I).** When compared to PV neurons of naïve mice, the spike count of PV neurons from nerve-injured mice was significantly lower (**Figure 3F, I**). However, after PV neurons of nerve injured mice were exposed to Lei-Dab7, this significant difference was lost, and we observed a partial recovery in the spike count (**Figure 3G, I**).

Our results demonstrate that PV neurons are equipped with the conductances required for frequency adaptation but block their activation presumably by preventing calcium accumulation. After nerve injury, these SK channels are activated, and induce frequency adaptation. Consequently, blocking these channels in PV neurons can restore their tonic firing.

### PGC-1α regulates PV expression in dorsal horn PV neurons

Nerve injury is associated with the activation of several spinal inflammatory pathways (Zhang and An 2007; Nadeau et al. 2011; Schomberg et al. 2012), some of which may presumably converge on the mechanisms regulating PV expression. Therefore, we sought to identify endogenous processes controlling PV expression in the spinal cord. In the cerebrum, one reported regulatory mechanism controlling the PV promoter activity is through a transcriptional coactivator named peroxisome proliferator-activated receptor-γ coactivator (PGC)-1α, referred to as PGC-1α (Lucas et al. 2010; Lucas et al. 2014). The expression of this coactivator is decreased by the TNFα pathway (Kim et al., 2007; Yang et al., 2017), and in the spinal cord, the expression of this inflammatory cytokine is increased after nerve injury (Gao et al., 2020). Thus, we first examined whether PGC-1α is expressed in dorsal horn PV neurons. Our fluorescence *in situ* hybridization results revealed that 80.5 ± 3.8% of neurons that co-express Pvalb and Ppargc1a (PGC-1α) also express Slc32a1 (VGAT) and are classified as inhibitory **(Figure 4A-C)**. Among the inhibitory PV neurons (that co-express Pvalb and Slc32a1), 65.8 ± 3.0% also expressed Ppargc1a above the detection threshold **(Figure 4D).** This data indicates that most dorsal horn inhibitory PV neurons co-express PGC-1α and that around 81% of PV neurons that express PGC-1α contain inhibitory markers.

**Figure 4.**
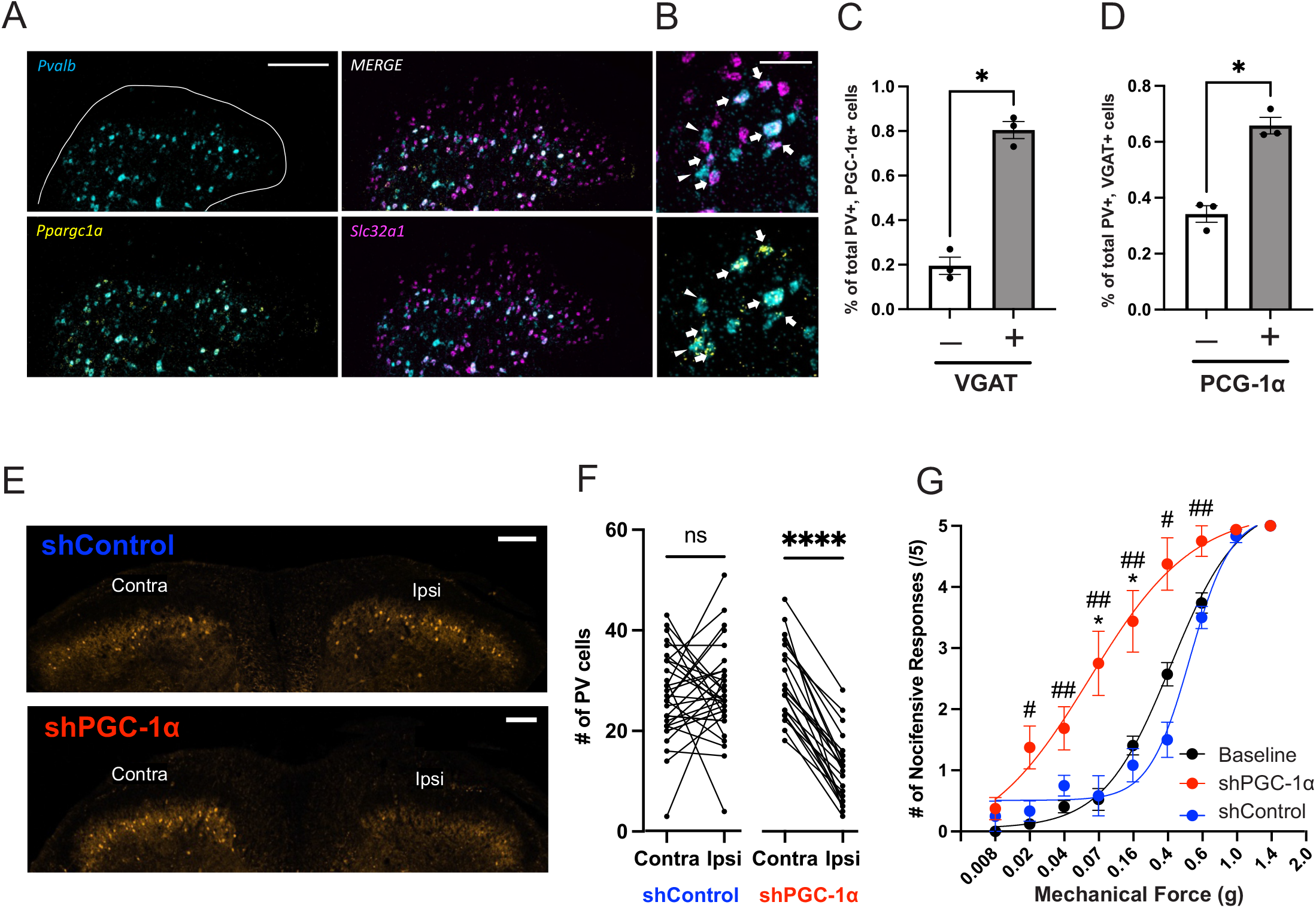
PGC-1α regulations PV expression in DH PV neurons. **A-B.** Fluorescence *in situ* hybridization of lumbar spinal dorsal horn sections showing PV mRNA (cyan), PGC-1α mRNA (ppgarc1a, yellow), and VGAT mRNA (slc32a1, magenta). Zoom in sections shows white arrows indicating VGAT+/PV+ cells colocalize with PGC-1α and white arrowheads show VGAT-/PV+ cells colocalize with PGC-1α. Scale bar, 100μm, 10μm. **C.** Mean (±SEM) of the percentage of PV+/PGC-1α+ cells with [VGAT+ (80.5 ± 3.8, 688/870 PV+/PGC-1α+ cells counted in 3 mice)] or without [VGAT-(19.5 ± 3.8, 182/870 PV+/PGC-1α+ cells counted)] VGAT. *p-value = 0.0156, paired t-test. (n=30 sections from 3 mice) **D.** Mean (±SEM) of the percentage of PV+/VGAT+ cells expressing PGC-1α [PGC-1α+ (65.8 ± 2.99, 688/1043 PV+/VGAT+ cells counted in 3 mice)] or not [PGC-1α-(34.2 ± 2.99, 355/1043 PV+/VGAT+ cells counted)]. *p-value = 0.0339, paired t-test. (n=30 sections from 3 mice) **E.** There is a decrease in tdTom+ cells (decrease in Pvalb promoter activity) on the ipsilateral side of injection of AD shPGC-1α (bottom) but no decrease in tdTom+ cells in the AD shControl condition (top) 4 days post injection. Scale bar, 200μm. **F.** Quantification of E showing no significant changes in the number of tdTom+ PV cells on the ipsilateral and contralateral side of AD shControl sections (n=32 sections in 3 mice). There is a significant decrease in the number of tdTom+ PV cells on the ipsilateral compared to contralateral side of AD shPGC-1α condition (n=22 sections in 2 mice). ****p-value <0.0001 paired t-test. **G.** Mean (±SEM) mechanically evoked (von Frey) nocifensive responses in shPGC-1α injected mice 4 days post injection (red dots and curve) show the development of mechanical allodynia. Mice injected with shControl (blue dots and curve) show no significant changes to their mechanical sensitivity compared to baseline (black dots and curve). Two-way ANOVA, Tukey’s multiple comparisons test, * compares shPGC-1α and shControl, # compares shPGC-1α and baseline. (n=8 shPGC-1α, n=6 shControl, n=14 baseline)

We then investigated whether PGC-1α regulates PV protein expression in PV neurons of the dorsal horn. We decreased PGC-1α expression with an intraspinal injection of an adenovirus carrying an shRNA against PGC-1α (shPGC-1α) in Pvalb-tdTomato mice. Our results indicate that mice treated with shPGC-1α showed a decrease in tdTomato+ cells on the side ipsilateral to the injection compared to the control virus (shControl) (**Figure 4E-F**) and compared to a spinal segment outside the injection site **(Figure S4)**. These results indicated that knocking down PGC-1α leads to a decrease in PV expression. Furthermore, when we tested the mechanical hypersensitivity of these mice four days post-injection, we observed the presence of mechanical allodynia in shPGC-1α-but not shControl-injected mice (**Figure 4G**). Our data suggest that a decrease in PGC-1α activity may be responsible for the decrease in PV expression after nerve injury.

### Increasing PVp expression prior to nerve injury prevents the development of mechanical allodynia

Our results have demonstrated that nerve injury produces a decrease in PV expression **(Figure 1P-R)**, and that this decrease is causally linked to the development of mechanical allodynia (F**igure 2**). Thus, we hypothesized that restoring PVp expression would prevent or alleviate nerve injury-induced mechanical allodynia. To test this hypothesis, we generated an AAV vector encoding a Cre-dependent HA-tagged PV (termed PV-WT), and a Cre-dependent inactive control in which PV lacked its two Ca^2+^ binding domains (termed PV-ΔEF, **Figure 5A**). We validated the function of these constructs in HEK293 cells stimulated with ATP (50μM, 4 sec), and observed that, unlike PV-ΔEF, PV-WT could prevent the rise in intracellular calcium (**Figure 5B**). To selectively target the expression of the HA-tagged constructs to dorsal horn PV neurons, we injected them in PV^Cre^;tdTom mice and confirmed their expression by immunohistochemistry experiments against the HA tag (**Figure 5C**). We then examined whether exogenously increasing PVp expression in PV neurons could prevent or reverse nerve injury induced mechanical allodynia. When we increased PVp expression two weeks before nerve injury, we observed (two weeks after nerve injury) that PV-WT injected mice developed significantly less mechanical allodynia compared to PV-ΔEF injected mice (**Figure 5D**). However, increasing PVp expression two weeks after nerve injury did not alleviate the mechanical allodynia (**Figure 5E**). Taken together, our results indicate that increasing PVp expression can prevent, but not alleviate, the development of mechanical allodynia after nerve injury.

**Figure 5.**
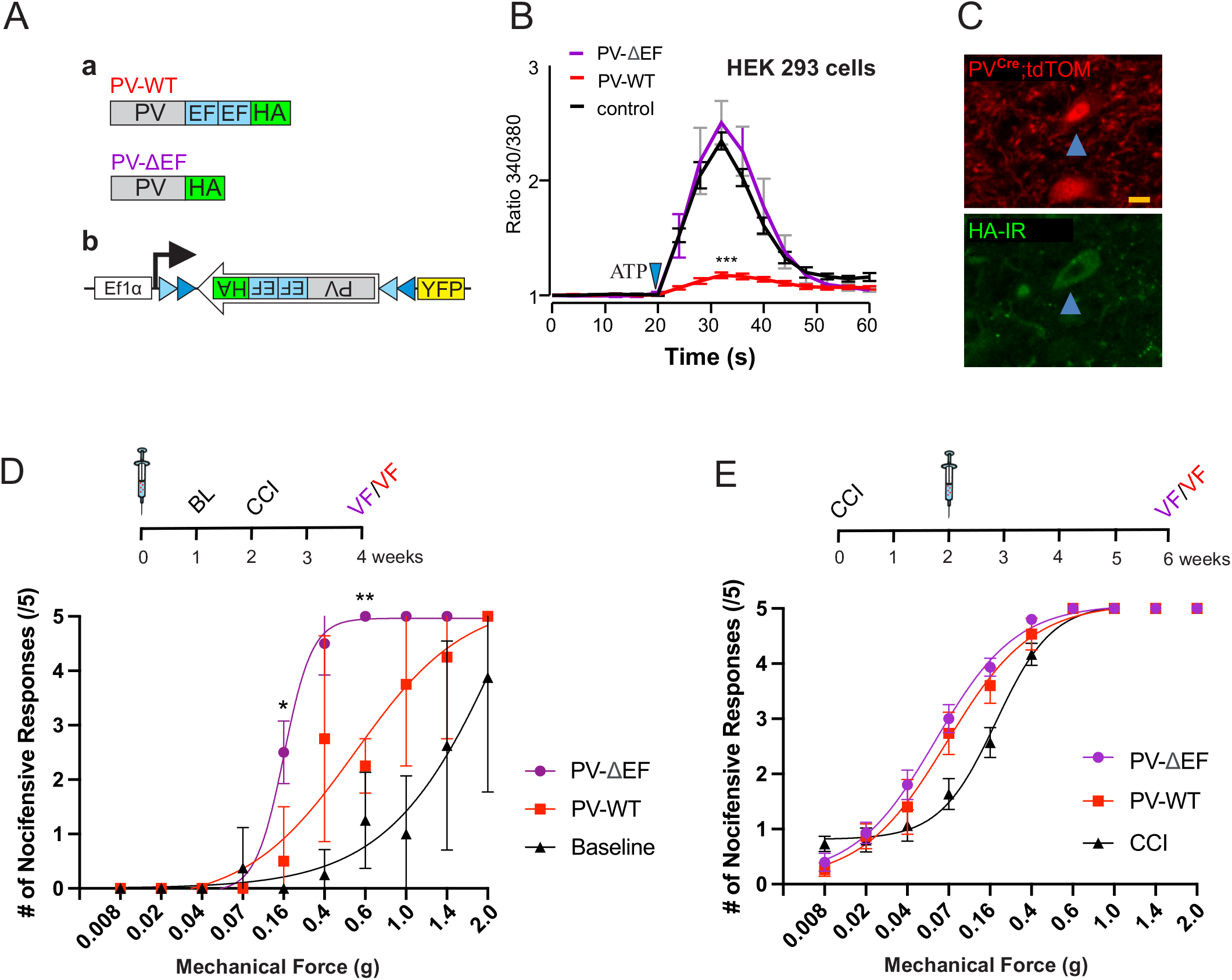
Re-establishing PVp in PV neurons prevents the development of mechanical pain after nerve injury. **Aa**. Plasmid construction of WT parvalbumin cDNA with its two EF calcium binding domains (PV-WT) and a truncated PV sequence lacking the EF domains (PV-ΔEF, bottom). Both sequences contain an HA-tag. **Ab**. Diagram depicting the construction of the AAV virus containing a YFP tag and the flexed PV-WT sequence to restore PVp expression only in PV^cre^ expressing neurons. **B.** Representative ATP-induced Ca^2+^ responses in HEK293 cells to functionally validate the plasmids construction. The rise in intracellular Ca^2+^ is significantly lower in the presence of PV-WT (red curve) but unaffected when PV-ΔEF is present (purple curve). The mean area under the curve from 20s to 50s was compared. (n= 369 cells PV-WT, n=277 PV-ΔEF, n= 35 control), One-way ANOVA **C.** Neuroanatomical section of PV^Cre^;tdTom neurons of the dorsal horn (red, top panel) that express (blue arrow) the PV-WT sequence revealed through its HA-tag (HA-IR; green, bottom panel). Scale bar, 10μm. **D.** Mean (±SEM) mechanically evoked (von Frey) nocifensive responses before (black triangles and curve) and after (red and purple traces) nerve injury in mice pre-injected with either the PV-ΔEF sequence (purple circles and curve) or the PV-WT sequence (red squares and curve). Two-way ANOVA, Tukey’s multiple comparisons test, PV-WT and baselines were not statistically significant. *compares PV-WT and PV-ΔEF. (n=4 PV-WT, n=4 PV-ΔEF). **E.** Mean (±SEM) mechanically evoked (von Frey) nocifensive responses of mice after nerve-injury treated with either the PV-ΔEF sequence (purple circles and curve) or the PV-WT sequence (red squares and curve) compared to CCI (black triangles and curve). No statistically significant changes in the mechanical hypersensitivity of PV-WT and PV-ΔEF. (n=5 PV-WT, n=5 PV-ΔEF) were observed.

## Discussion

Our results established a pivotal role for the calcium binding protein parvalbumin in the development of mechanical hypersensitivity. The model we propose is one in which the decrease in PV expression after nerve injury leads to an accumulation in intracellular Ca^2+^, activating SK channels and transitioning the neuron to adaptive firing (**Figure 6**).

**Figure 6.**
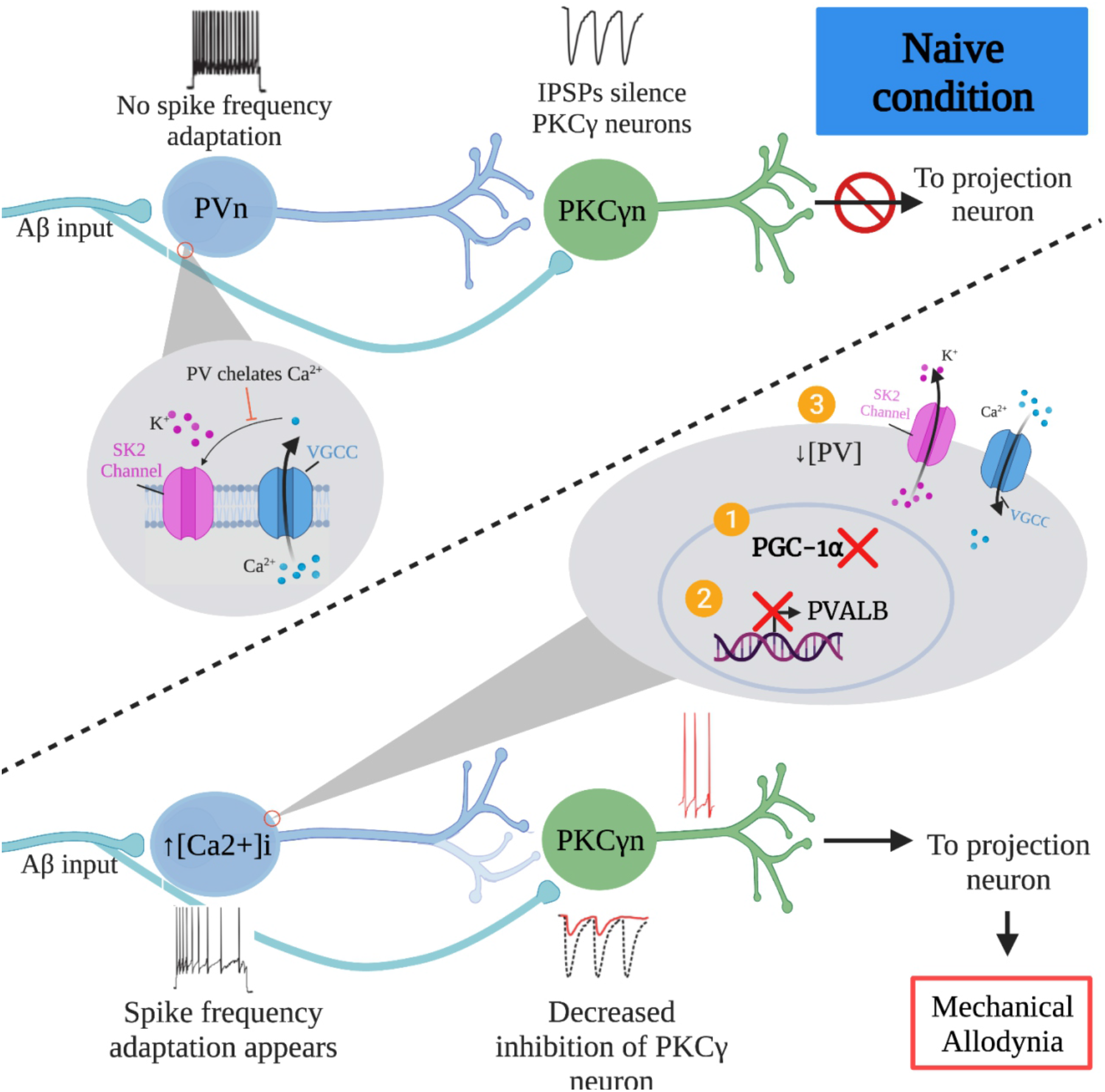
General organization of PV neurons and PKCγ neurons relationship in normal and neuropathic pain conditions. Under naive conditions (top panel), PV protein in the PV neurons chelates the free intracellular calcium ions, thereby inhibiting the activation of SK2 channels (magenta in the grey circle, top half) and preventing the efflux of K+ ions. No PV neuron spike frequency adaption occurs, thus allowing for inhibition of the polysynaptic targets and subsequent activation of lamina I projection neurons. After nerve injury (bottom half), we propose that a decrease in PGC-1α (1) elicits a decrease in PV expression (2), resulting in an accumulation in free intracellular calcium and activation of SK2 channels (3) that allow for spike frequency adaptation of PV neurons. This adaptation decreases their inhibition of PKCγ neurons, enabling low-threshold inputs to activate the excitatory neurons and the subsequent activation of lamina I projection neurons to cause mechanical allodynia.

Although in most CNS regions PV-expressing neurons are inhibitory, the dorsal horn represents a unique area in which the calcium buffer is also expressed in excitatory neurons (Häring et al., 2018; Sathyamurthy et al., 2018, Russ et al., 2021). This feature is not restricted to mice, as recent data from rhesus macaque (Arokiaraj et al., 2022) and human (Yadav et al., 2022) dorsal horns indicate that PV is expressed in both glutamatergic and GABAergic neuronal clusters. The percentage of excitatory and inhibitory neurons expressing PV has been studied in mice and is less consistent, varying from 50 to 95% inhibitory (Hughes et al., 2012; Petitjean et al., 2015; Abraira et al., 2017; Gradwell et al., 2022). These disparities can be due to the different PV^Cre^ driver lines used to identify PV neurons (Jax stock #008069; or #017320). Another source of variability is in the markers used in immunohistochemistry (VGAT vs. Pax2) or *in situ* (VGAT vs. Gad1) experiments, or in the image analysis parameters used in *in situ* hybridization experiments. At the level of the mRNA, we found around 65% of Pvalb expressing cells were inhibitory (Slc32a1 probe) with the reminder excitatory (Slc17a6 probe). Interestingly, we observed the presence of a small portion (<10%) of PV neurons that expressed similar intensities of Slc17a6 and Slc32a1 mRNAs. These neurons were classified as excitatory or inhibitory based on the marker that was expressed at higher intensity (**Figure S1C**). It is important to note that these observations do not consider any post-translational regulatory mechanisms. Ultimately, the clinical relevance of these rodent models remains inconclusive without insights into the organization of the human spinal cord.

Recent advances in ISH and IHC on human spinal cord tissue (Shiers et al., 2021; Yadav et al., 2022; Arokiaraj et al., 2022; Dedek et al., 2022) provide the needed resources that will help validate our preclinical findings of inhibitory neurons. In the mouse, the dorsal horn is comprised of several subsets of inhibitory interneurons involved in modality-specific processing of peripheral inputs (Bourane et al., 2015a; Bourane et al., 2015b; Koch et al., 2017). Multiple reports have highlighted the importance of the inhibitory tone provided by PV neurons (Hughes et al., 2012; Petitjean et al., 2015; Abraira et al., 2017; Cao, Scherrer & Chen, 2022), and modeled their impact on the excitability of dorsal horn circuits (Medlock et al., 2022). Our results suggest that the decrease in PV expression leads to the decrease in PV neuron output. However, this chronological sequence of events has been shown to be different in other CNS areas. For instance, it was reported that in the hippocampus, enhanced photoactivation of PV neurons elevated the proportion of PV neurons with high expression of the calcium buffer (Donato, Rompani & Caroni, 2013). More recently, Kourdougli and colleagues demonstrated that chronic (4 days) pharmacological activation of PV precursor neurons (using Nkx2.1^Cre^ mice) caused an increase in PV expression (Kourdougli et al., 2022). Although our work clearly demonstrates a correlation between PVp expression and the inhibitory output of these neurons, the chronological order of these events remains to be determined.

Overall, the loss of inhibitory output from PV neurons after peripheral nerve injury could be due to at least three mechanisms. *The efficacy of their synaptic connection* with their targets could be reduced, through pruning of their terminals, decreased vesicular content of their inhibitory transmitters, or decreased post-synaptic receptors. This hypothesis is supported by a recent study demonstrating that PV neurons lose their inhibitory effect on postsynaptic targets after nerve injury (Cao, Scherrer & Chen, 2022). The mechanisms that cause this reduced inhibition remain unclear. One explanation is the pruning of PV synapses to their postsynaptic targets. Indeed, we have shown a decrease in the number of appositions between PV neurons and PKCγ somata after spared nerve injury (SNI) leaving afferents from the tibial nerve intact (Petitjean et al., 2015). However, another report using a SNI model sparing the sural afferents did not observe a decrease in the number of inhibitory PV synapses onto the cell body and dendrites of PKCγ neurons (Boyle et al., 2019). This discrepancy between the two models could be due to topographical differences in the termination of these sciatic nerve branches within the mediolateral zones of the dorsal horn (Corder et al., 2010). Although the role of PKCγ in nerve injury-mediated mechanical allodynia is established (Malmberg et al., 1989; Petitjean et al., 2015; Miraucourt, Dallel & Voisin, 2007), as well as their inputs from PV neurons (Petitjean et al., 2015; Boyle et al., 2019; Cao, Scherrer & Chen, 2022), it is possible that the decrease in PV-PKCγ synapses alone may not be the only explanation for the reduced inhibitory output from PV neurons.

A second possibility could be the *decreased excitatory drive onto PV neurons* from peripheral Aβ fibers. It was reported that after nerve injury, the amplitude of Aβ-fiber stimulated excitatory postsynaptic potentials recorded in lamina III glycinergic neurons was decreased and failed to generate action potentials (Lu et al., 2013). Despite being unidentified, these recorded inhibitory neurons were described to have islet cell-like morphology and tonic firing patterns, characteristics of inhibitory PV neurons. However, a more recent study observed no significant change in the number of Aβ boutons onto the processes of inhibitory PV neurons (Boyle et al., 2019), suggesting that if there is a decreased excitatory drive onto PV neurons, it is the result of weakened synaptic function rather than structural integrity. Together, these studies suggest an increase in failure rate of synaptic transmission between Aβ and inhibitory PV neurons after peripheral nerve injury.

A third possibility for the reduced inhibitory output of PV neurons after nerve injury is a *change in their intrinsic electrical properties*. We speculated that if a reduced Aβ input alone was responsible for the decreased activity of PV neurons after nerve injury, we should be able to directly stimulate the soma of these inhibitory neurons and observe a firing pattern similar to naïve mice. However, our results indicate that PV neurons from nerve injured mice respond differently to the injection of depolarizing currents. Indeed, after nerve injury, these neurons display not only a reduced number of action potentials in response to current injections, as seen by us and others (Boyle et al, 2019), but also a change in their firing pattern. We observed the presence of spike frequency adaptation, suggesting that nerve injury recruits calcium-activated potassium conductances. We demonstrated that the adaptive firing observed in PV neurons of nerve injured mice could be partially reversed by blocking SK2 channels, confirming their role in this phenomenon and that additional mechanisms are also at play. The increased contribution of these potassium channels is presumably the result of impaired calcium buffering caused by the decrease in PVp expression after nerve injury. In fact, we demonstrate that reducing PVp expression alone in an otherwise healthy animal was sufficient to elicit mechanical allodynia and change the firing activity of PV neurons.

It is unclear the extent to which each of these three mechanisms contribute to the decreased output of dorsal horn PV neurons and which mechanism should be targeted for therapeutic pain relief. Our study showed that rescuing PV protein alone after nerve injury is not sufficient to alleviate mechanical allodynia. Further work is required to investigate the signaling pathways upstream of the regulation of PV expression in order to modulate its expression level after nerve injury.

In conclusion, we propose a model in which the incoming inputs from Aβ fibers are prevented from polysynaptically activating lamina I projection neurons because PV neurons receive a copy of this input and can impose it on their post-synaptic targets, whether they are PKCγ neurons (Petitjean et al., 2015; Cao, Scherrer & Chen, 2022; Medlock et al, 2022), vertical cells (Boyle et al., 2019), or the central terminals of Aβ fibers themselves (Hughes et al., 2012; Abraira et al., 2017; Boyle et al., 2019). Such forward inhibitory mechanism is only possible if the firing pattern of PV neurons can match exactly the continuous firing pattern of the Aβ fibers. If the PV neurons develop frequency adaptation, they will not efficiently prevent the activation of their targets, which will initiate the polysynaptic activation of lamina I projection neurons involved in pain signaling. Approaches aimed at increasing the excitability of PV neurons may therefore have therapeutic potential and should thus be further investigated.

## Materials and Methods

### Mouse lines and animal care

Male mice (8-14 weeks old) were kept on a 12-h:12-h light/dark cycle, with food and water provided ad libitum. All experimental procedures were approved by the Animal Care and Use Committee at McGill University, in accordance with the regulations of the Canadian Council on Animal Care. All mouse strains have previously been described. PV^Cre^, B6 PV^Cre^, Pvalb-tdTomato, PV^Cre^-tdTom reporter, Ai65F (RCF tdTom) and Pv-CreERT2 (PV KO) were obtained from The Jackson Laboratory (Catalog #008069, #017320, #027395, #007914, #032864, and #010777 respectively). C57BL/6 mice aged 8-12 weeks were purchased from Charles Rivers.

### Molecular Biology

Mice were anesthetized with 5% isofluorane and perfused transcardially with ice-cold oxygenated (95% O2, 5% CO2) N-Methyl-D-glutamine-based artificial cerebrospinal fluid (NMDG-ACSF) solution (compounds previously outlined under electrophysiology). The lumbar spinal segment was rapidly removed and immersed in ice-cold NMDG-ACSF and the spinal cord dorsal ipsilateral and contralateral sections were dissected free. The spinal cord tissue was frozen on dry ice and stored at –80° C. Total RNA was extracted using Relia Prep Tissue RNA system (Promega, Z6111) and converted to cDNA using SuperScript™ IV VILO™ Master Mix (Thermo Fisher Scientific – Invitrogen, 11756050). For each sample, qPCR reactions were performed in triplicates on Applied Biosystems Step One Plus Real Time qPCR System using TaqMan FastAdvanced Master Mix (Thermo Fisher Scientific - Applied Biosystems, 4444557) and Taqman Gene expression Assays (Thermo Fisher Scientific – Applied Biosystems) for the following genes: Kcnn1 (Assay ID: Mm01349167_m1), Kcnnn2 (Assay ID: Mm00446514_m1), Kcnn3 (Assay ID: Mm00446516_m1), Kcnn4 (Assay ID: Mm00464586_m1), Parvalbumin (Assay ID: Mm00443100_m1), Ppargc1a (Assay ID: Mm01208835_m1), PKC gamma (Assay ID: Mm00440861_m1), AIF1 (Assay ID: Mm00479862_g1) and GFAP (Assay ID: Mm01253033_m1). Gene expression levels were quantified by the double delta Ct analysis using Beta Actin (Assay ID: Mm00607939_s1) or Tbp (Assay ID: Mm00446973_m1) as housekeeping reference genes. The housekeeping reference genes were selected based on results from spinal cord samples of 2 naïve and 2 CCI mice obtained from TaqMan™ Array Mouse Endogenous Controls Plates, Fast 96-well (Thermo Fisher Scientific – Applied Biosystems, 4426694) and can be found in Supplementary Table 1. For each experiment, we selected the control gene whose threshold cycle value was closest to our gene of interest.

### RNAscope In Situ Hybridization

Mouse lumbar spinal cords were extracted following an intraperitoneal injection of 2,2,2-Tribromoethanol (Avertin, 250mg/kg) for deep anesthesia. Then, the tissue was embedded in Optimal Cutting Temperature compound (OCT) and flash frozen on dry ice. Transverse sections (14 mm) were cut using the cryostat (Lecia Microsystems), directly mounted on Fisherbrand Superfrost Plus Microscope Slides (Cat. No. 12-550-15) and stored on dry ice before starting the RNAscope in situ hybridization. Stainings were acquired using Advanced Cell Diagnostics RNAscope® Multiplex Fluorescent Reagent Kit V2 (Cat. No. 323100): a TSA-based Fluorescent RNAscope®detection assay with 4-plex capability (enabled by the addition of RNAscope 4-Plex Ancillary Kit for Multiplex Fluorescent v2, Cat. No. 323120) used in combination with Opal fluorophores from Akoya Biosciences. Briefly, slides of fresh frozen spinal cord sections from WT mice were fixed for 15 minutes in cold 4% PFA (prepared in 1xPBS). After fixation, the sections were dehydrated in ethanol and pretreated with Hydrogen Peroxide and Protease IV (Cat. No. 322381) at room temperature for 30 minutes. Pretreated sections were incubated for 2 hours at 40C in ACD HybEZ oven (Cat. No. 241000ACD) with RNAscope probes against VGAT (Probe - Mm-Slc32a1-C2, reference 319191-C2), Pvalb (Probe-Mm Pvalb-C3, reference 421931-C3) and VGlut2 (Probe - Mm-Slc17a6-C4, reference 319171-C4), PGC-1α (Probe – Mm-Ppargc1a-C1, reference 402121), SK2 (Probe-Mm-Kcnn2-C2, reference 427971-C2), SK3 (Probe-Mm-Kcnn3-C4, reference 427961-C4). Control slides were run simultaneously and treated with a mouse RNAscope® 4-Plex Positive Control Probe-Mm (reference 321811) and RNAscope® 4-Plex Negative Control Probe (reference 321831). The signal was amplified through a series of three hybridization steps using amplification reagents included in the RNAscope® Multiplex Fluorescent Reagent Kit v2. The signal was revealed by HRP using the following Probe-Opal dye combinations: Akoya Biosciences Opal™ 520 Reagent Pack (Cat. No.FP1487001KT); Opal™ 570 Reagent Pack (Cat. No. FP1488001KT); Opal™ 620 Reagent Pack (Cat. No. FP1495001KT) and Opal™ 690 Reagent Pack (Cat. No. FP1497001KT). Finally, the sections were stained with DAPI and mounted using ProLong Gold antifade reagent (Invitrogen, Cat. No. P36930).

### Tissue preparation and Immunohistochemistry

Immunohistochemistry was performed as previously described (Petitjean et al., 2019). Briefly, mice were anesthetized with 5% isofluorane and perfused transcardially with 0.1 M saline phosphate buffer (PBS; [in mM] 154 NaCl, 13 Na2HPO4, 2.5 NaH2PO4, pH 7.4) followed by with 4% paraformaldehyde (PFA) in PBS. Spinal cords were then extracted by laminectomy, postfixed for 2 hours in same fixative, and cryoprotected in 30% sucrose / PBS solution at 4°C. Lumbar spinal cord sections were cut using a cryostat (Leica Microsystems). Transverse sections (25 μm-thick) were cut and placed in a 48-well plate containing PBS and stored at 4°C. Tissue sections were washed three times in PBS/0.3% Triton X-100 (PBS-T; Sigma, St. Louis, MO), and incubated in 10% normal goat serum/PBS/0.3% Triton X-100 (NGST) for 1 hour, before adding the primary antibodies over 48 hours at 4°C in a 1% NGST solution. Sections were then washed three times with PBS-T solution and incubated for 1 hour with Alexa fluorophore-conjugated secondary antibodies at room temperature (RT). The sections were washed three times with PBS, air-dried, and coverslipped using Aqua Polymont (Polysciences, Inc., Warrington, USA). The slides were stored protected from light at 4°C until analysis.

Primary antibodies were used in 1% NGST at the indicated concentrations: anti-parvalbumin rabbit (1:1000; Swant PV27), anti-Iba1 rabbit (1:1000, Wako #019-19741), anti-Pax2 goat (1:1000, R&D Systems #AF3364), anti-Tlx3 rabbit (1:1000, Abcam #1840111), anti-HA mouse (1:1000, Cedarlane #MMS-101P), anti-WGA rabbit (1:1000, Sigma-Aldrich #T4144), anti-PKCγ guinea pig (1:1000, gift from Allan Basbaum). All primary staining were revealed with corresponding secondary antibodies conjugated to Alexa fluorophores, diluted at 1:500 in 1% NGST. All sections were incubated with NucBlue Fixed Cell Stain (DAPI) for 5 minutes at RT before mounting on microscope slides.

### Microscopy and image quantification

Images for quantitative analysis were acquired using either a Zeiss LSM780 or a Zeiss LSM710 scanning confocal microscope. All imaging was performed on 20x/0.40LD Plan-Neofluar objective lens at 1024X1024 pixels/frame, 0.6 digital zoom and 12-bit using a 32 GaAsp detecter array. The lasers were: laser diode 405nm 30mW, Ar Ion laser 458/488/514nm 25mW, DPSS-laser 561nm 20mW, HeNe Green 543nm 1mW, HeNe 594nm 2mW, HeNe RED 633nm 5mW. Image files were processed for analysis in ImageJ (NIH) using the “Blind Analysis Tools” plug-in by two independent observers. For RNAscope quantification, an intensity threshold for each probe was set at 4 times the mode to define individual cells as positive or negative for mRNA expression.

### Chronic Constriction Injury Surgery

Unilateral Chronic constriction injury of the sciatic nerve (CCI) was done as previously described (Austin, Wu & Moalem-Taylor, 2012). Three ligatures (6-0 Perma-Hand silk suture; Ethicon) were placed around the sciatic nerve bundle and loosely closed to construct but not arrest epineural blood flow. Mice developed robust mechanical allodynia that becomes chronic after two weeks and can persist for up to 8 weeks post-surgery.

### AAV virus production

HEK293T cells were co-transfected with pHelper, pAAV2/8 and pAAV-EF1a-DIO-msParvalbumin-IRES-NLS-EYFP or pAAV-EF1a-DIO-msParvalbumin Δ 13-37-IRES-NLS-EYFP. On the third day post-transfection, cells were harvested and lysed. The virus containing supernatant was collected and purified by iodixanol density gradient ultracentrifugation, followed by concentration with Amicon Ultra-15 10K centrifugation filtres. Titration was performed by qPCR and measured in GC/ml.

### Intraspinal viral injections

This method was described in a previous study (Petitjean et al., 2019). Briefly, mice were anesthetized with 5% isoflurane/oxygen and maintained with 2.5% during the operation. A 1.5 cm incision was made on the back of the animal to expose the underlying spinal column at the lumbar level. Muscle and connective tissues overlaying L4-L5 spinal segments were removed, and the spinal column was immobilized in a stereotaxic frame. A glass microelectrode was lowered through the dura mater at the intervertebral space between the T13 and L1 vertebral segments (AP 0, ML 0.45, DV 0.25 mm) and 300 nL of virus was injected (50 nl/min). The electrode was slowly removed after 2.5 min. Carprofen analgesic (5 mg/kg) was given 20 min prior to the beginning of the surgery and every 24 h for 3 days after the surgery. Mice recovered on a heat pad before being returned to their cage. Mice continued to be group housed.

The viruses were AAV2/2-EF1α-DIO-WGA (Titer: 3.44E11 GC/ml), Lenti PLKO.1-puro-CMV-TGFP-si/shRNA PV (Canadian Neurophotonics Platform, Titer: 4.5E9 TU/mL), Lenti-PLKO.1-puro-CMV-TGFP-non target control (Canadian Neurophotonics Platform, Titer: 12E10 TU/mL), AAV2/8-EF1a-DIO-msPV-ires-NLS-EYFP, AAV2/8-EF1a-DIO-msPVΔ-ires-NLS-EYFP. The AD shPGC-1α (shPGC-1α: GGTGGATTGAAGTGGTGTAGA, Titer: 1.16E10 ifu/mL) and AD shControl (shControl: ACAACAGCCACAACGTCTATA, Titer: 4.32E10 ifu/mL) viruses were gifted from Dr. Estall (Besse-Patin et al., 2019). Both constructs were generated using the AdEasy system. The vector is AdTrack with a U6 promoter driving the shRNA.

### von Frey assay

Mechanical sensitivity was assessed as previously described (Petitjean et al., 2015). Mechanical withdrawal threshold was assessed by placing mice on an elevated wire-mesh grid and stimulating the plantar surface of the hindpaw with von Frey filaments. Starting with a low weight filament, that exerted 40 mg of pressure, each filament was applied 5 times against the hindpaw. If no withdrawal response was elicited after the 5 applications, the next filament (stronger) was applied and so on until 5 of 5 applications elicited a response. The filament where 5 of 5 applications elicited a response was deemed the mechanical threshold. Animals were tested 3 times, once every day before the spinal cord injection of the AAVs. Before slice electrophysiology of CCI mice, we verified their development of mechanical allodynia by testing mechanical sensitivity before and 2-4 weeks post CCI.

The PKCγ antagonist was dissolved in physiological saline and injected at a concentration of 100 pmoles in a volume of 5 μl in the intrathecal space of isoflurane-anesthetized mice. The PKCγ blocker γV5-3 is derived from amino acids 659-664 of PKCγ conjugated to TAT peptide (residues 47-57) by an S-S bond via Cys residues added to the N terminus of each of the peptides. The TAT peptide enables entry of γV5-3 into cells. Consequently, the unconjugated TAT peptide was used as control (in 5 μl saline).

### Radiant heat paw-withdrawal assay

The withdrawal latency to high temperature was measured using the radiant heat paw-withdrawal (Hargreaves) test (Stoelting). Mice were placed into individual restrainers on a glass platform 15 min before the experiment. A noxious thermal stimulus was focused through the glass onto the plantar surface of a hindpaw until the animal withdrew the paw from the heat source. The paw-withdrawal latency was automatically measured to the nearest 0.1 s. A cut-off latency of 20 s was used to avoid tissue damage.

### Calcium imaging

During calcium imaging experiments, HEK293 cultured cells were continuously superfused with an extracellular saline solution containing: 130 mM NaCl, 5 mM KCl, 2.5 mM CaCl2,1mM MgCl2, 10 mM glucose, and 10 mM Hepes (pH 7.4). The superfusion rate was 1.5 mL/min for a total bath volume of 1 mL. Cells were loaded with the calcium-sensitive dye Fura-2 by incubating the cultures for 1 h at room temperature (22–25 °C) in the extracellular solution containing 2 mM Fura-2 acetoxymethyl ester and 0.001% (w/v) pluronic acid. Cells were washed three times with extracellular saline solution before and after loading. Fluorescence measurements on individual cells were monitored using a NeuroCCD-SM256 imaging system (RedShirtImaging, Decatur, GA) mounted on a Zeiss Axio Examiner microscope with filter set 46HE and a 40x water immersion objective (0.75 NA). Images were acquired at 50 Hz with a 103 digital gain. Image analysis was performed by drawing a region of interest around individual cells. Cells were alternately excited at wavelengths of 340 nm and 380 nm with a filter wheel, and emitted light was collected above 520 nm. Pairs of images were acquired every 2 s. Intracellular calcium level and their variations are expressed as the ratio of fluorescence signals (ratio F340/F380) measured at 520 nm after alternate excitation at 340 nm and 380 nm. This ratio was calculated after background signal subtraction. All experiments were performed at room temperature (22–25 °C).

### Electrophysiology

Parasagittal lumbar spinal cord slices were obtained from adult male mice (12 to 16 weeks old). Animals were deeply anesthetized through an intraperitoneal injection of 2,2,2-Tribromoethanol (Avertin, 250mg/kg). Mice were then briefly perfused transcardially with an ice-cold oxygenated (95% O2, 5% CO2) N-Methyl-D-glucamine-based artificial cerebrospinal fluid (NMDG-ACSF) solution containing the following (in mM): 93 NMDG, 93 HCl 12M, 2.5 KCl, 1.25 NaH2PO4, 30 NaHCO3, 20 HEPES, 25 glucose, 2 thiourea, 5 Na-L-ascorbate, 3 Na-pyruvate, 10 N-acetyl-L-cysteine, 0.5 CaCl2/2H2O, and 10 MgSO4/7H2O (pH 7.3–7.4). The lumbar spinal segment was rapidly removed and immersed in ice cold NMDG-ACSF and the spinal cord was dissected free. The ventral roots were removed and 300 μm transverse slices with attached dorsal roots were cut with a vibratome (Leica VT1000S). Slices were transferred to a submerged chamber containing HEPES-based recovery ACSF for 10 minutes at 34 C, equilibrated with 95% O2 and 5% CO2. Following the recovery incubation, slices were transferred to a recording chamber and continuously superfused with oxygenated ACSF (in mM): 126 mM NaCl, 26 mM NaHCO3, 2.5 mM KCl, 1.25 mM NaH2PO4, 2 mM CaCl2, 2 mMMgCl2, and 10 mM glucose (bubbled with 95% O2 and 5% CO2; pH 7.3; 310 mOsm measured), where they were then maintained at room temperature prior to transfer to the recording chamber.

### Patch clamp

Slices were transferred to a recording chamber and continuously superfused with oxygenated ACSF (32-34 C, 2 ml/min). Patch pipettes were pulled from borosilicate glass capillaries (Harvard Apparatus) with a P-97 puller (Sutter Instruments). They were filled with a solution containing 80 mM K2SO4, 10 mM HEPES, 2 mM KCl and 2 mM MgCl2 (pH 7.3, adjusted with CsOH; osmolarity, 310 mOsm, adjusted with sucrose) and had final tip resistances of 6–8 MU for whole-cell recording. Pipettes were filled with a solution containing 145 mM NaCl, 5 mM KCl 10 mM HEPES, 1 mM MgCl2 and 1 mM CaCl2 (pH 7.3, adjusted with CsOH; osmolarity, 300 mOsm, adjusted with sucrose) and had final tip resistances of 6–8 megaohm for recordings. Neurons were viewed by an upright microscope (Olympus) with a 40X water-immersion objective, infrared differential interference contrast (IR-DIC) and fluorescence. Recordings were made from identified PV-neurons expressing the tdTomato. Data were acquired with pClamp 10.0 software (Molecular Devices) using MultiClamp 700B patch-clamp amplifier and Digidata 1440A (Molecular Devices). Recordings were low-pass filtered on-line at 2 kHz, digitized at 20 kHz.

### Illustrations

All figures presented in this manuscript were assembled CorelDRAW Graphic Suite. Graphics were created with BioRENDER (https://biorender.com/).

### Statistics

All statistical analyses were performed using GraphPad Prism v9.3.0 software and presented as mean ± standard error of the mean (SEM). A p-value of p < 0.05 was the criteria for significance.

## Acknowledgments

Images were collected for this manuscript was performed in the McGill University Advanced BioImaging Facility (ABIF), (RRID:SCR_017697) and IRCM Microscopy and Imaging Platform. We are thankful to Ms. T. Koch at the McGill University Comparative Medicine and Animal Resources Centre for the mice colony management. We thank Dr. M. Millecamps and the McGill Animal Behavioural Characterization platform for the mouse behaviour experimental support.

