## Supplementary Materials for "Parvalbumin protein controls inhibitory tone in the spinal cord"

**Supplementary Table 1.** *Endogenous control genes cycle thresholds ($C_T$) of naïve and CCI dorsal horn*

Endogenous control genes used in this study were selected based on the $C_T$ of the genes of interest. Pes1, Rps17, and Tfrc have different cycle thresholds and are not recommended to be used as controls when comparing various genes in naïve and CCI (highlighted in yellow).

**Figure S1**. *Co-expression of PV mRNA with VGAT or VGLUT2 mRNA*

**A.** The intensity of the VGAT (Slc32a1) probe and the VGLUT2 (Slc17a6) probe in Pvalb+ cells classified as inhibitory (left) or excitatory (right).

**B.** The VGAT intensity (purple) and the VGLUT2 (blue) intensity of each Pvalb+ cell. Cells in the purple box were classified as inhibitory because they had a greater intensity of Slc32a1 and cells in the blue box were classified as excitatory because they had a greater intensity of Slc17a6, with the vertical black line separating the two populations. Around 10% of the total Pvalb+ cells had similar expression of VGAT and VGLUT2 mRNA (< 2 folds difference).

**C.** Zoomed in image of dorsal horn *in situ* hybridization. White arrows show colocalization of Pvalb and Slc17a6 indicating excitatory PV interneurons. White arrowhead shows colocalization of Pvalb and Slc32a1 indicating inhibitory PV interneurons. White asterisk (*) shows Pvalb neuron that expresses both Slc17a6 and Slc32a1. NucBlue is shown in grey. Scale bar, 50 µm.

**D-E.** The specificity and sensitivity of $PV^{cre}$;tdTom (red) mice through labeling parvalbumin-immunoreactive neurons (green) in the lumbar dorsal horn. Scale bar, 100µm. On the graph, dots represent individual measurements and lines represent mean ±SEM. (n=26 sections in 3 mice)

**F.** Transverse lumbar spinal sections of the dorsal horn showed increased microglia activation (Iba1) in CCI mice compared to naïve. Scale bar, 100µm.

**G.** There is an increased ipsilateral/contralateral ratio of Iba1 intensity in the dorsal horn of CCI mice compared to naive. ****p-value <0.0001 unpaired t-test.

**H.** Histogram of individual tdTomato+ cell intensity of CCI (red) and naïve (blue) mice.

**Figure S2.** *Decreasing PV in the dorsal horn elicits mechanical allodynia*

**A-B.** Transverse dorsal horn spinal cord sections of Pvalb-tdTomato mice that received intraspinal delivery of a lentivirus carrying shRNA against PV or NT and the ratio of tdTomato+ cells 8 weeks post injection (69.14 ± 6.4% n=24 sections from 3 mice injected with shPV; 91.61 ± 0.3% n=24 sections from 3 mice injected with NT). *p-value=0.0364, unpaired t-test

**C.** Quantification of the paw withdrawal latency in the Hargreaves' test. There is no difference between the latencies of mice injected with shRNA against PV (shPV, n=5) or shRNA non-targeting (NT, n=5) before the injection (baseline, BL) or after 1-, 4-, and 8-weeks post injection (W1, W4, W8). Two-way ANOVA, Šídák's multiple comparisons test.

**D.** Change in mechanical sensitivity after intraspinal injection of a lentivirus carrying shRNA against PV (n =5, red circles and curve) shows the development of mechanical allodynia. Mice receiving non-targeting shRNA (n =5, black squares, and curve) are unaffected. Two-way ANOVA, *p-value=0.0192, Šídák's multiple comparisons test comparing shPV and NT. #p-value=0.0108, ##p-value=0.0013, Šídák's multiple comparisons test comparing shPV and Baseline.

**E.** Left dorsal horn spinal injection of an AAV carrying the WGA transsynaptic tracer in a double inverted orientation (AAV2/2-EF1α-DIO-WGA), is first only expressed by tdTomato+ PV neurons (red, box i). Then PV neurons transfer WGA to their post synaptic targets (blue)

which do not colocalize with $PV^{cre}$;tdTom neurons. Some of these post synaptic targets are PKCγ expressing neurons (PKCγ, green, box ii). Scale bar, 10μm.

**F**. Quantification of the neurons that contain WGA in the ipsilateral dorsal horn of the injection site. WGA+ neurons either colocalize with $PV^{cre}$;tdTom neurons (yellow, 10 out of 284 neurons examined), or PKCγ+ neurons (magenta, 50 out of 284 neurons examined), or neither (blue, 224 out of 284 neurons). No colocalization of WGA+ neurons with PKCγ+/PV neurons were found on the contralateral side of the same section (n = 11 sections from 3 mice).

**G.** Mean (±SEM) mechanically evoked (von Frey) nocifensive response threshold as a percent of baseline response in $PV^{cre}$;RCF tdTomato mice injected with shRNA-Pvalb (30.1 ± 7.4%, n= 7 mice, red bar) or non-targeting (52.79 ± 4.3%, n=7 mice, grey bar).

**Figure S3** *Tonic firing persists after HCN channel blockade*

**A-D.** Whole cell current clamp raw traces illustrating the response to incremental current step injections in a PV neuron recorded in lumbar spinal cord slices of a naïve mouse before and after application of HCN channel blocker, ZD3288 (10μM). The spike count analysis **(C)** and resting membrane potential **(D)** (mean ± S.E.M, n=5 cells from n=2 naïve mice) are shown. *p<0.01 Two-way ANOVA, Šídák's multiple comparisons test. n.s. non-statistically significant, paired t-test

**Figure S4** *PGC-1α controls PV promoter activity*

Outside the AD shPGC-1α injection site (green box), there is no decrease in the number of tdTom+ PV cells on the ipsilateral compared to contralateral side.

**Supplementary Table 1**

| Gene | Naive | CCI |
|---|---|---|
| 18s | 10.108514 | 9.670838 |
| Abl1 | 23.899994 | 23.88978 |
| Actb | 18.891298 | 18.921028 |
| ATP5 | 16.861061 | 16.981333 |
| B2m | 21.819077 | 21.871689 |
| Casc3 | 30.903704 | 30.922707 |
| Cdkn1a | 24.889587 | 24.905165 |
| Cdkn1b | 25.90512 | 25.888596 |
| Eif2b1 | 24.91807 | 24.929529 |
| Elf1 | 25.830839 | 25.90757 |
| Gadd45a | 25.88781 | 25.858072 |
| Gapdh | 17.869818 | 17.907259 |
| Gusb | 25.750452 | 25.705687 |
| Hmbs | 26.902388 | 26.913261 |
| Hprt1 | 22.826517 | 22.840378 |
| Ipo8 | 24.909086 | 24.880728 |
| Mrpl19 | 24.943712 | 24.940247 |
| Pes1 | 25.92412 | 26.963074 |
| Pgk1 | 20.874527 | 20.93853 |
| polr2a | 22.883829 | 22.888628 |
| Pop4 | 25.973955 | 25.934288 |
| Ppla | 18.875296 | 18.872744 |
| Psmc4 | 23.918831 | 23.935837 |
| Pum1 | 23.917353 | 23.942045 |
| Rpl30 | 26.909744 | 26.807648 |
| Rpl37a | 23.913389 | 23.876074 |
| Rplp2 | 30.900206 | 30.944944 |
| Rps17 | 18.198736 | 20.30422 |
| Tbp | 25.948351 | 25.916662 |
| Tfrc | 22.940729 | 23.925318 |
| Ubc | 19.898195 | 19.928846 |
| Ywhaz | 20.850203 | 20.86817 |

### Figure S1

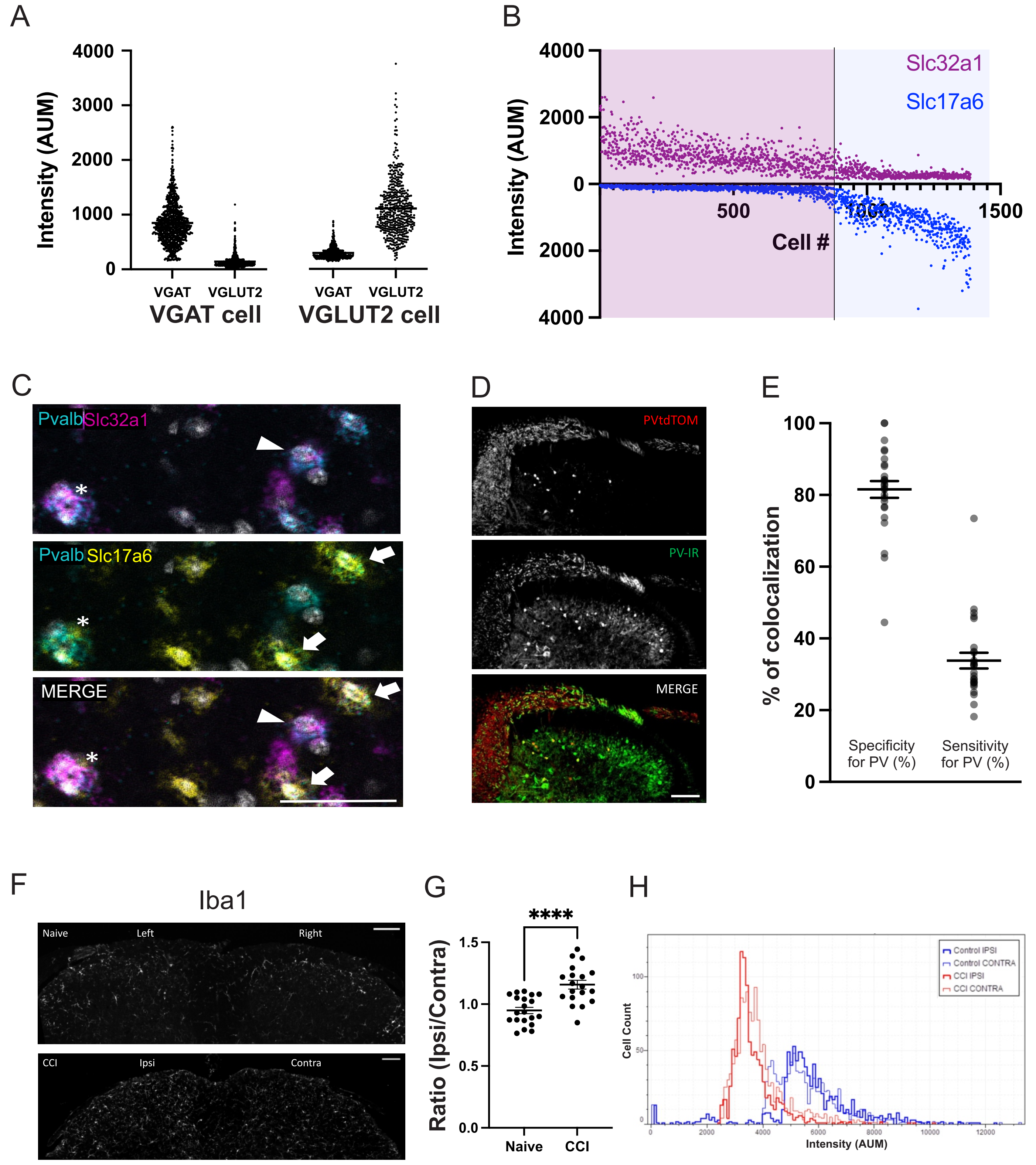

A
Intensity (AUM)
4000
3000
2000
1000
0
VGAT
VGLUT2
VGAT
VGLUT2
VGAT cell
VGLUT2 cell
B
Slc32a1
Slc17a6
Intensity (AUM)
4000
2000
0
2000
4000
500
1000
1500
Cell #
C
Pvalb Slc32a1
Pvalb Slc17a6
MERGE
D
PVtdTOM
PV-IR
MERGE
E
% of colocalization
100
80
60
40
20
0
Specificity for PV (%)
Sensitivity for PV (%)
F
Iba1
Naive
Left
Right
CCI
Ipsi
Contra
G
****
Ratio (Ipsi/Contra)
1.5
1.0
0.5
0.0
Naive
CCI
H
Control IPSI
Control CONTRA
CCI IPSI
CCI CONTRA
Cell Count
Intensity (AUM)

### Figure S2

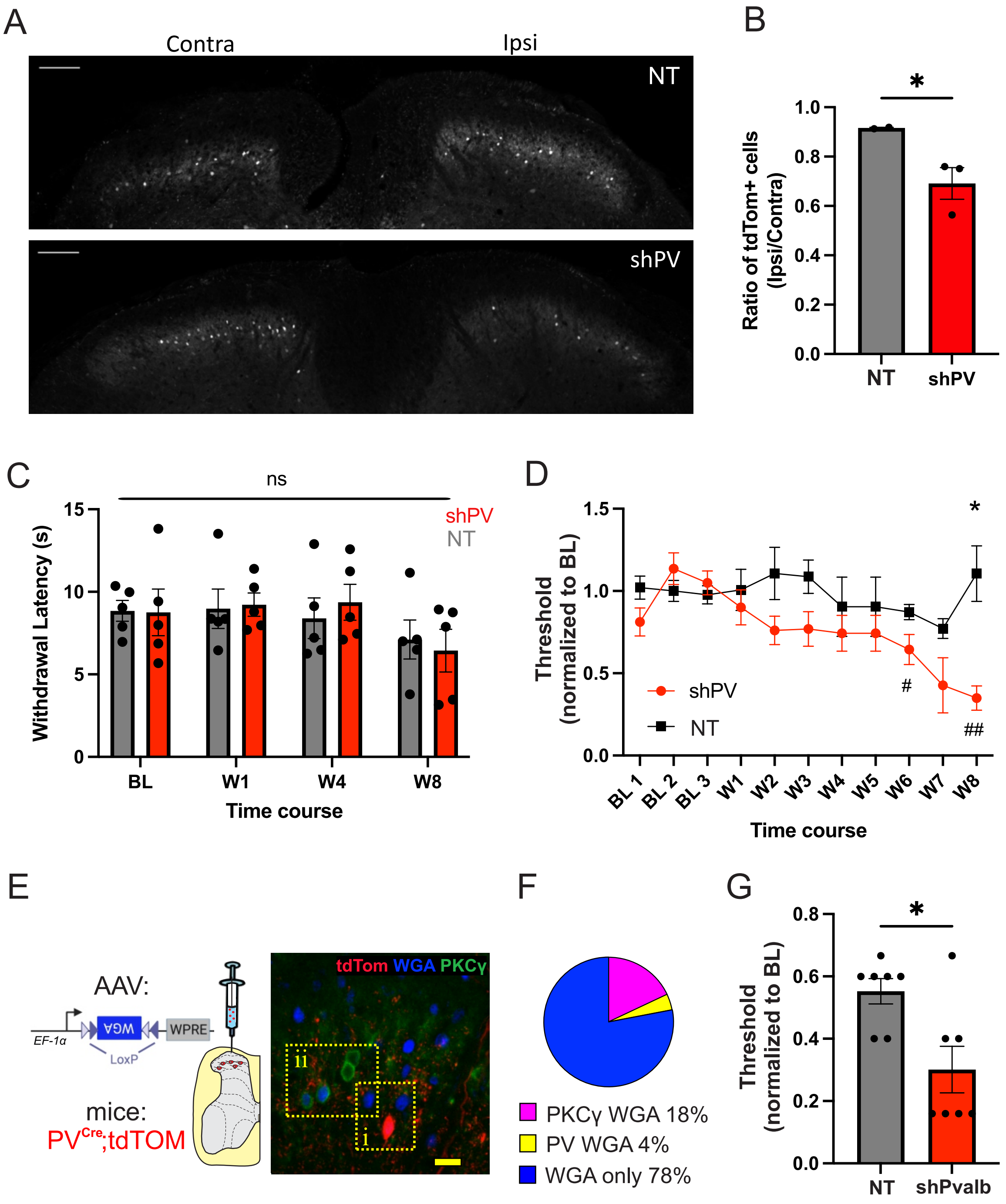

A
Contra
Ipsi
NT
shPV
B
*
Ratio of tdTom+ cells (Ipsi/Contra)
1.0
0.8
0.6
0.4
0.2
0.0
NT
shPV
C
ns
shPV
NT
Withdrawal Latency (s)
15
10
5
0
BL
W1
W4
W8
Time course
D
*
Threshold (normalized to BL)
1.5
1.0
0.5
0.0
shPV
NT
#
##
BL 1
BL 2
BL 3
W1
W2
W3
W4
W5
W6
W7
W8
Time course
E
AAV:
EF-1α
WGA
WPRE
LoxP
mice:
PVCre;tdTOM
tdTom WGA PKCγ
ii
i
F
PKCγ WGA 18%
PV WGA 4%
WGA only 78%
G
*
Threshold (normalized to BL)
0.8
0.6
0.4
0.2
0.0
NT
shPvalb

### Figure S3

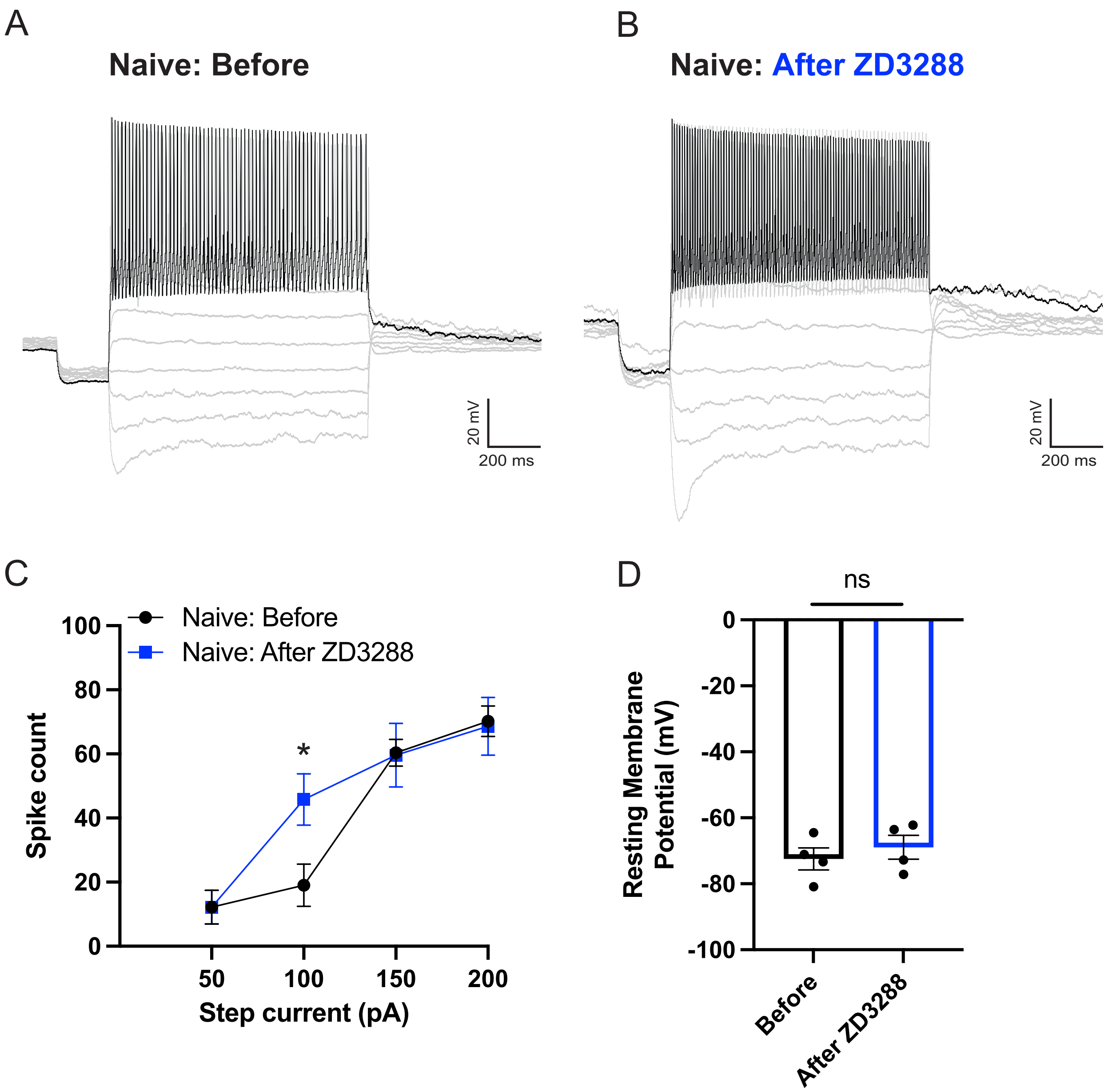

A
Naive: Before
20 mV
200 ms
B
Naive: After ZD3288
20 mV
200 ms
C
Naive: Before
Naive: After ZD3288
Spike count
100
80
60
40
20
0
*
50
100
150
200
Step current (pA)
D
ns
Resting Membrane Potential (mV)
0
-20
-40
-60
-80
-100
Before
After ZD3288

### Figure S4

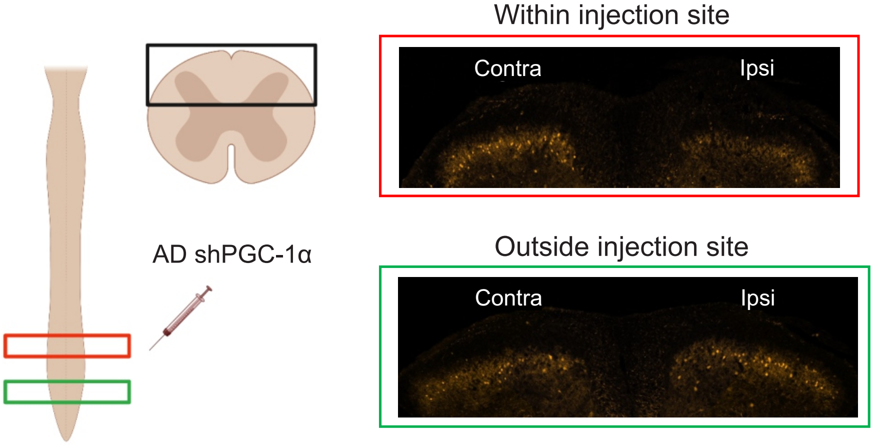
AD shPGC-1α
Within injection site
Contra
Ipsi
Outside injection site
Contra
Ipsi
